# Robust identification of perturbed cell types in single-cell RNA-seq data

**DOI:** 10.1101/2023.05.06.539326

**Authors:** Phillip B. Nicol, Danielle Paulson, Gege Qian, X. Shirley Liu, Rafael Irizarry, Avinash D. Sahu

## Abstract

Single-cell transcriptomics has emerged as a powerful tool for understanding how different cells contribute to disease progression by identifying cell types that change across diseases or conditions. However, detecting changing cell types is challenging due to individual-to-individual and cohort-to-cohort variability and naive approaches based on current computational tools lead to false positive findings. To address this, we propose a computational tool, *scDist*, based on a mixed-effects model that provides a statistically rigorous and computationally efficient approach for detecting transcriptomic differences. By accurately recapitulating known immune cell relationships and mitigating false positives induced by individual and cohort variation, we demonstrate that *scDist* outperforms current methods in both simulated and real datasets, even with limited sample sizes. Through the analysis of COVID-19 and immunotherapy datasets, *scDist* uncovers transcriptomic perturbations in dendritic cells, plasmacytoid dendritic cells, and FCER1G+NK cells, that provide new insights into disease mechanisms and treatment responses. As single-cell datasets continue to expand, our faster and statistically rigorous method offers a robust and versatile tool for a wide range of research and clinical applications, enabling the investigation of cellular perturbations with implications for human health and disease.

## Introduction

The advent of single-cell technologies has enabled measuring transcriptomic profiles at single-cell resolution, paving the way for the identification of subsets of cells with transcriptomic profiles that differ across conditions. These cutting-edge technologies empower researchers and clinicians to study human cell types impacted by drug treatments, infections like SARS-CoV-2, or diseases like cancer. To conduct such studies, scientists must compare single-cell RNA-seq (scRNA-seq) data between two or more groups or conditions, such as infected versus non-infected (1), responders versus non-responders to treatment (2), or treatment versus control in controlled experiments.

Two related but distinct classes of approaches exist for comparing conditions in single-cell data: differential abundance prediction and inter-condition difference identification. Differential abundance approaches, such as DA-seq, Milo, and Meld (3; 4; 5; 6), focus on identifying cell types with varying proportions between conditions. In contrast, inter-condition difference identification methods, the focus of this study, seek to detect cell types with distinct transcriptomic profiles between conditions (7; 8). It is essential to distinguish between these two classes of problems, as their outputs cannot be compared directly.

Past studies have mostly relied on manual approaches involving visually inspecting data summaries to detect differences in scRNA data (7). Specifically, cells were clustered based on gene expression data and visualized using *uniform manifold approximation* (UMAP). Cell types that appeared *separated* between the two conditions were identified as different. To avoid depending on a manual approach, *Augur*, a computational method specifically developed to detect cell-types that differ across conditions, employed a machine learning technique to quantify the cell-type specific separation between groups (8). However, none of the current ad-hoc or computational approaches take into consideration individual-to-individual variation. Note that cells are more similar within individuals than across individuals (9). This variation is why methods that perform statistical inference, such as *limma, edgeR*, and *DESeq2* (10; 11; 12) are widely used in bulk gene expression studies. Because this source of variation is partially natural (driven by genetics, for example), changing the technology used to measure expression does not eliminate it (13). Furthermore, in scRNA-Seq experiments, individual-to-individual variability can also be introduced by technical artifacts, for example, if samples from different individuals are sequenced at different times or collected from different sites.

In this study, we show that by failing to account for individual-to-individual variability, current approaches can incorrectly identify cell-type-specific between-group differences. To address this, we developed a statistical approach that models individual-to-individual and technical variability explicitly and provides a way to quantitatively define transcriptomic differences between groups. The approach, which we call *scDist*, is based on a linear mixed-effects model of single-cell gene expression counts. Furthermore, because transcriptomic profiles are high-dimensional, we develop an approximation for the between-group differences, based on a low-dimensional embedding, which results in a computationally convenient implementation that is substantially faster than *Augur*. We demonstrate the benefits using a COVID-19 dataset, showing that *scDist* can recover biologically relevant between-group differences while also controlling for sample-level variability. Furthermore, we demonstrated the utility of the *scDist* by jointly inferring information from five single-cell immunotherapy cohorts, revealing significant differences in a subpopulation of NK cells between immunotherapy responders and non-responders, which we validate in bulk transcriptomes from 789 patients. These results highlight the importance of accounting for individual-to-individual and technical variability for robust inference from single-cell data.

## Results

### Not accounting for individual-to-individual variability leads to false positives

We used blood scRNA-seq from six healthy controls (1) (see **Table 1**), and randomly divided them into two groups of three, generating a negative control dataset in which no cell type should be detected as being different. We then applied *Augur* to these data. This procedure was repeated 20 times. *Augur* falsely identified several cell types as perturbed (**Fig 1a**). *Augur* quantifies differences between conditions with an area under the curve (AUC) summary statistic, related to the amount of transcriptional separation between the two groups (AUC = 0.5 represents no difference). Across the 20 negative control repeats, 93% the AUCs > 0.5 and red blood cells (RBCs) were identified as perturbed in all 20 trials (**Fig 1a**). This false positive result was in part due to high across-individual variability in cell types such as RBCs (**Fig 1b**).

**Table 1:** Datasets used in the figures. The small COVID-19 dataset was generated by (1) and can be downloaded as a Seurat object from www.covid19cellatlas.org. The simulated null data can be generated using the *scDist* R package. The large COVID-19 was generated by (18) and can be accessed from the respective publication. Immunotherapy cohorts used in Figure 5 perform integrative analysis of 4 scRNA datasets for discovery, and for validatation, 7 bulk-RNA-Seq datasets are utilized.

| Figure | Dataset |
| --- | --- |
| Figure 1 | Small COVID-19 (1), Simulated null data |
| Figure 3 | Small COVID-19 (1), Simulated null data |
| Figure 4 | Large COVID-19 (18) |
| Figure 5 | Immunotherapy cohorts :<br>• 4 scRNA datasets (discovery) (24; 2; 25; 26)<br>• bulk-RNA-Seq datasets (validation) (33; 34; 35; 36; 37; 38; 39) |

**Figure 1:**
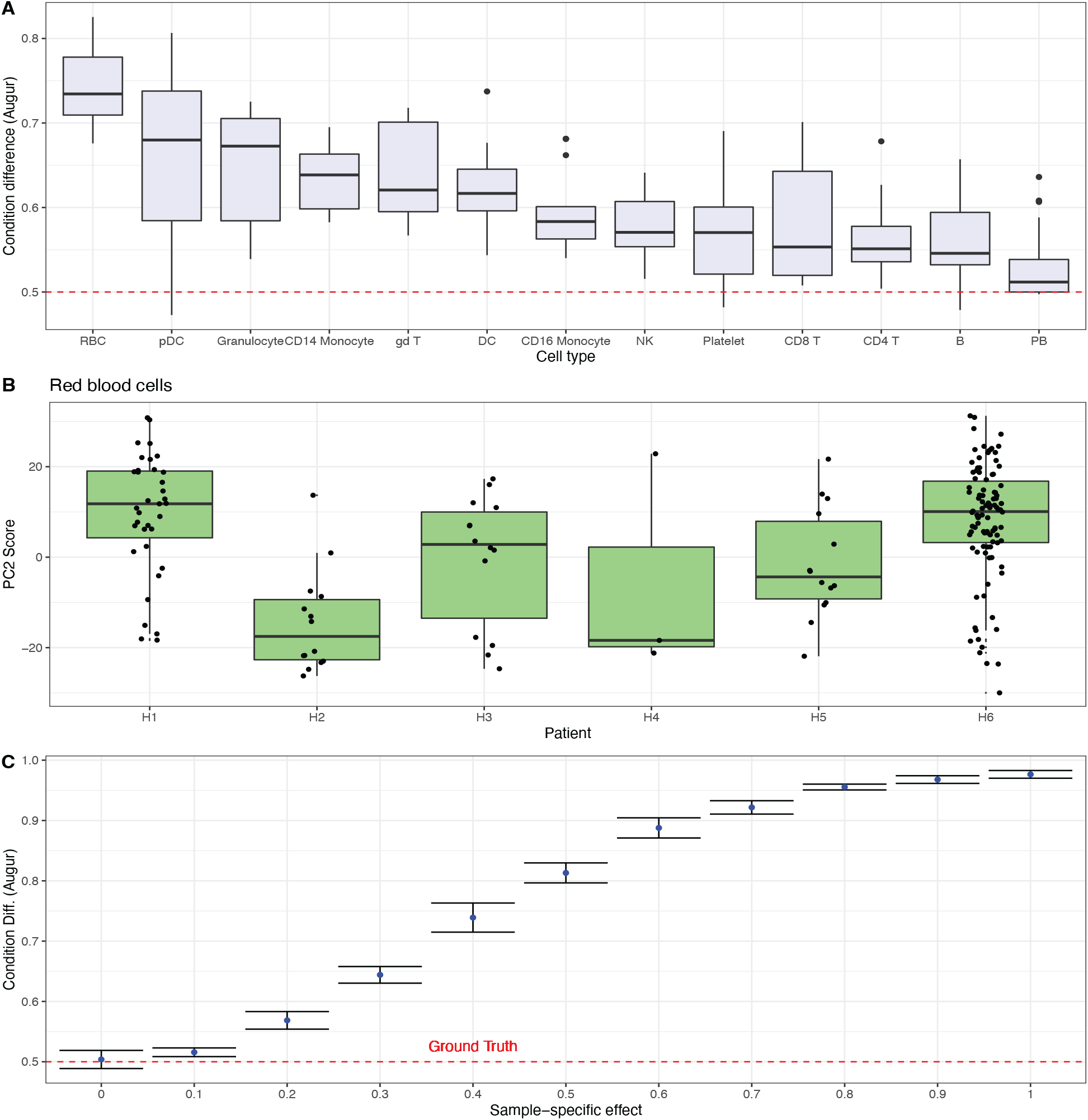
Evaluating *Augur* ‘s Performance in Negative Control Experiments. **A)** AUCs achieved by *Augur* on 20 random partitions of healthy controls, with no expected cell type differences (dashed red line indicates the null value of 0.5). **B)** Boxplot depicting the second PC score for red blood cells from healthy individuals, highlighthing high across-individual variability (each box represents a different individual). **C)** AUCs achieved by *Augur* on simulated scRNA-seq data with no condition differences but varying patientlevel variability (dashed red line indicates the ground truth value of no condition difference, AUC 0.5), illustrating the influence of individual-to-individual variability on false positive predictions.

We confirmed that individual-to-individual variation underlies false positive predictions made by *Augur* using a simulation. We generated simulated scRNA-seq data with no condition-level difference and varying patient-level variability (Methods). As patient-level variability increased, differences estimated by *Augur* also increased, converging to the maximum possible AUC of 1 (**Fig 1c**): *Augur* falsely interpreted individual-to-individual variability as differences between conditions.

*Augur* recommends that unwanted variability should be removed in a pre-processing step using batch correction software. We applied Harmony (14) to the same dataset (1), treating each patient as a batch. We then applied *Augur* to the resulting batch corrected PC scores and found that several cell types still had AUCs significantly above the null value of 0.5 (**Fig S1a**). On simulated data, batch correction as a pre-processing step also leads to confounding individual-to-individual variability as condition difference (**Fig S1b**).

### A mixed-effect model controls for false positives

To account for individual-to-individual variability, we modeled the vector of normalized counts with a linear mixed-effects model. Specifically, for a given cell type, let *z*_*ij*_ be a length *G* vector of normalized counts for cell *i* and sample *j* (*G* is the number of genes). We then model

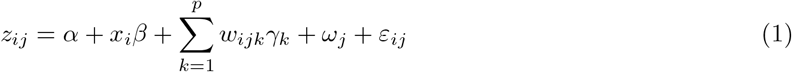

where *α* is a vector with entries *α*_*g*_ representing the baseline expression for gene *g, x*_*j*_ is a binary indicator that is 0 if individual *j* is in the reference condition, and 1 if in the alternative condition, *β* is a vector with entries *β*_*g*_ representing the difference between condition means for gene *g, w*_*ijk*_ are additional covariates that can encode batches or individual characteristics, the *γ*_*k*_ are vectors of effects related to the covariates *w*_*ijk*_, *ω*_*j*_ is a random effect that represents the differences between individuals, and *ε*_*ij*_ is a random vector (of length *G*) that accounts for other sources of variability. We assume that the *ω*_*j*_ are independent of each other and *ε*_*ij*_ follows a normal distribution with mean 0 and covariance matrix *τ* ^2^*I*. We assume that the *ε*_*ij*_ are independent of each other and also follow a normal distribution with expected value 0 and covariance matrix *σ*^2^*I*.

To obtain normalized counts, we recommend defining *z*_*ij*_ to be the vector of Pearson residuals obtained from fitting a Poisson or negative binomial GLM (15), the normalization procedure is implemented in the scTransform function (16). However, our proposed approach can be used with other normalization methods for which the model is appropriate.

Note that in model (1), the expected values for the two conditions are *α* and *α* + *β*, respectively, and that the more non-zero entries *β* has, the more different the conditions are. Therefore, we quantified the difference in expression profiles with the 2-norm of the vector *β*:

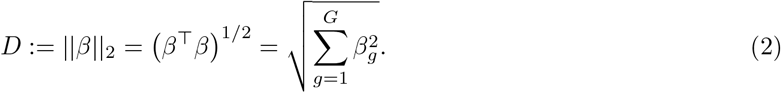

Here, *D* can be interpreted as the Euclidean distance between condition means (**Fig 2a**).

**Figure 2:**
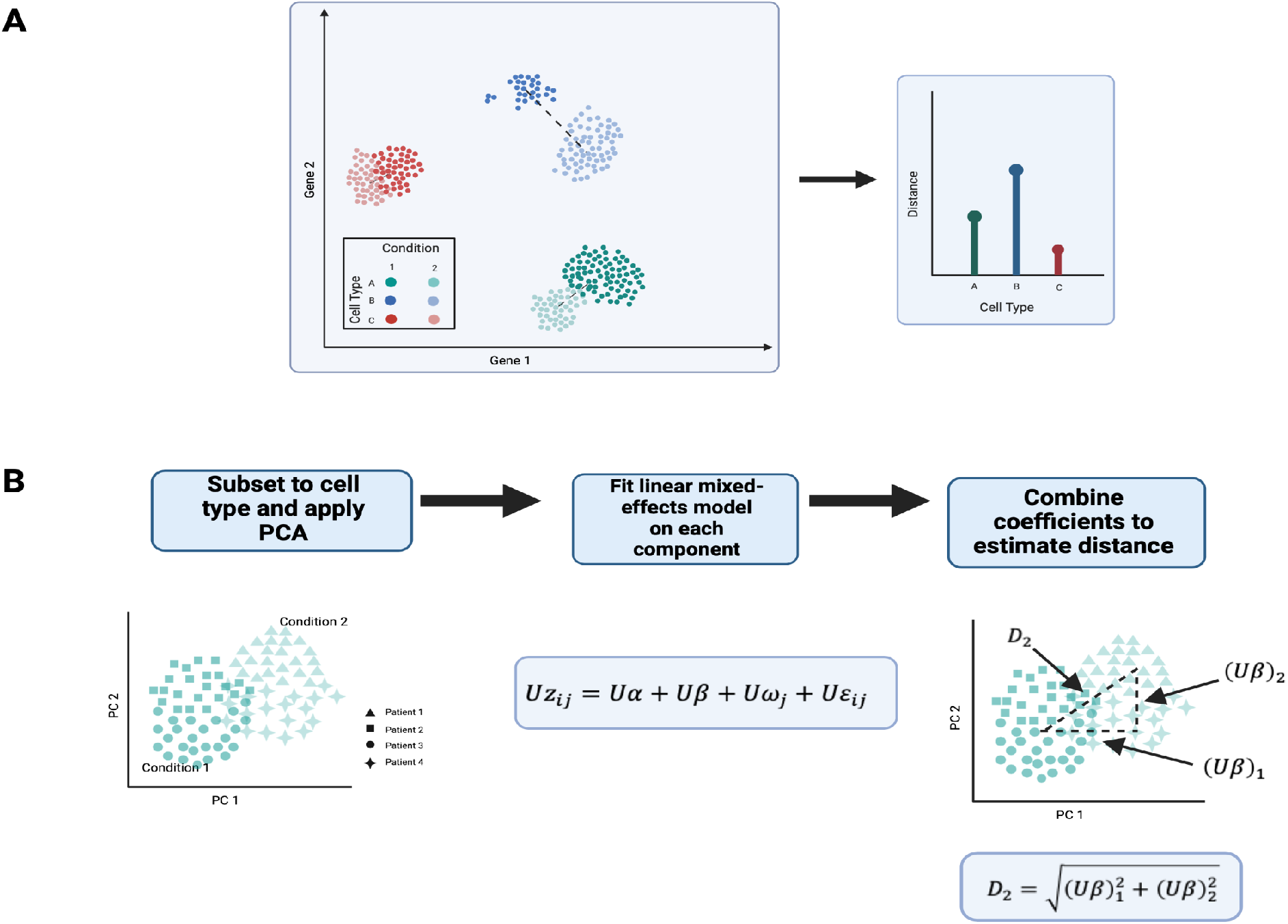
Visual Representation of the *scDist* Method. **A)** *scDist* estimates the distance between condition means in high-dimensional gene expression space for each cell type. **B)** To improve efficiency, *scDist* calculates the distance in a low-dimensional embedding space (derived from PCA) and employs a linear mixed-effects model to account for sample-level and other technical variability.

Because we expected the vector of condition differences *β* to be sparse, we improved computational efficiency by approximating *D* with a singular value decomposition (SVD) to find a *K* × *G* matrix *U*, with *K* much smaller than *G*, and

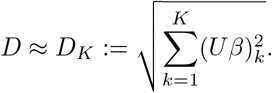

With this approximation in place, we fitted model equation (1) by replacing *z*_*ij*_ with *Uz*_*ij*_ to obtain estimates of (*Uβ*)_*k*_. A challenge with estimating *D*_*K*_ is that the maximum likelihood estimator can have significant upward bias when the number of patients is small (as is typically the case). For this reason, we employed a post-hoc Bayesian procedure to shrink 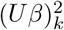 towards zero and compute a posterior distribution of *D*_*K*_ (17). We also provided a statistical test for the null hypothesis that *D*_*K*_ = 0. We refer to the resulting procedure as *scDist* (**Fig 2b**). Technical details are provided in Methods.

We applied *scDist* to the negative control dataset based on blood scRNA-seq from six healthy used to show the large number of false positives reported by *Augur* (**Fig 1**) and found that the false positive rate was controlled (**Fig 3a**). We then applied *scDist* to the data from the simulation study and found that, unlike *Augur*, the resulting distance estimate does not grow with individual-to-individual variability (**Fig 3b**). *scDist* also accurately estimated distances on fully simulated data (**Fig S2**).

**Figure 3:**
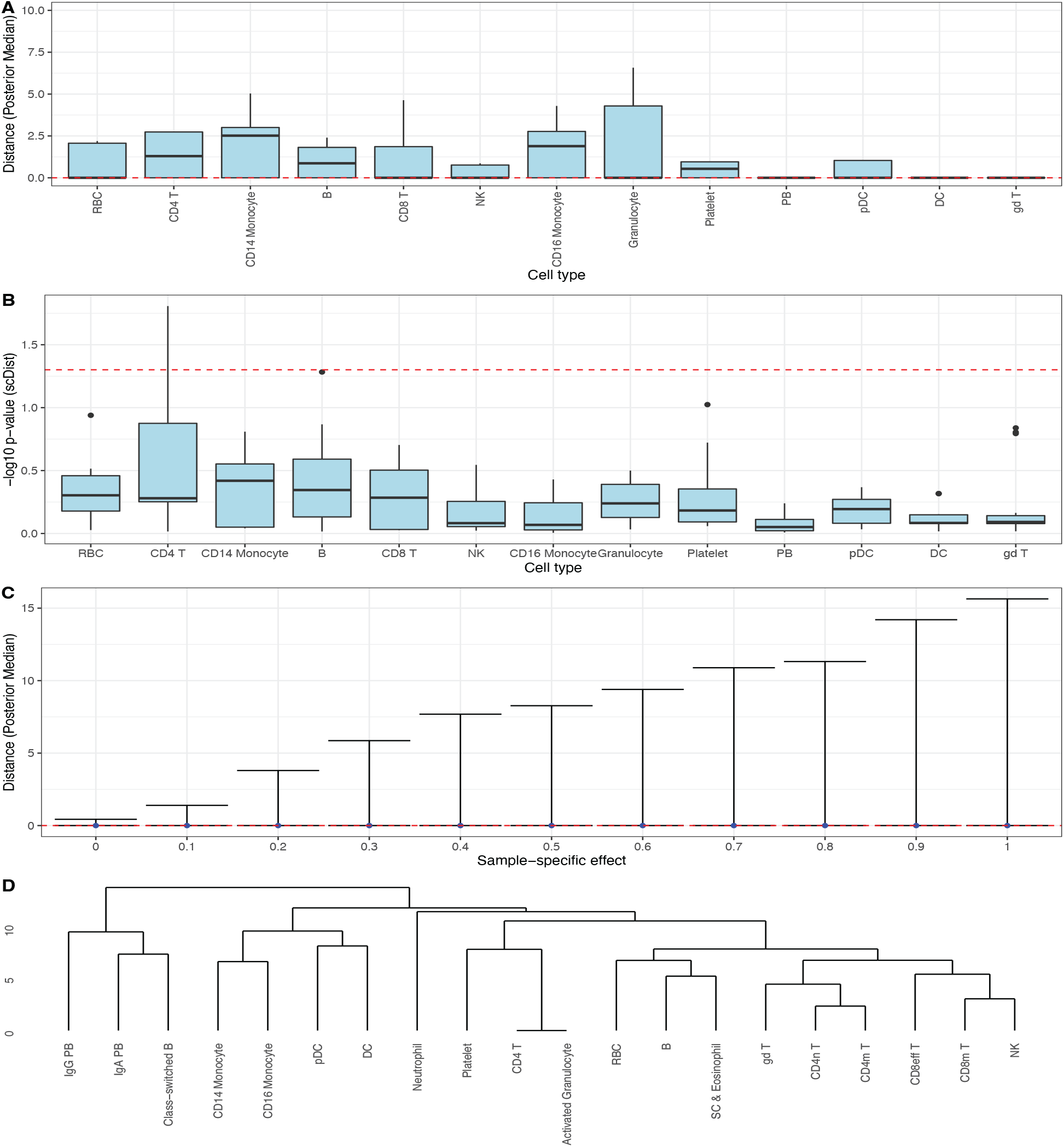
Application and Performance of *scDist*. **A)** Reanalysis of the data from **Fig 1b** using distances calculated with *scDist* ; the dashed red line represents ground truth value of 0. **B)** As in **A)**, but for p-values computed with *scDist* ; values above the dashed red line represent *p <* 0.05. **C)** Null simulation from **Fig 1c** reanalyzed using distances calculated by *scDist* ; the dashed red line represents the ground truth value of 0. **D)** Dendrogram generated by hierarchical clustering based on distances between pairs of cell types estimated by *scDist*.

The above results demonstrated that accounting for individual-to-individual variability greatly improves specificity over *Augur*. To determine if the improvement in specificity was achieved without loss of sensitivity, we applied *scDist* to obtain estimated distances between pairs of known cell types in the COVID-19 patients dataset (1) and then applied hierarchical clustering to these distances. The resulting clustering is consistent with known relationships driven by cell lineages (**Fig 3c**). Specifically, Lymphoid cell types T and NK cells clustered together, while B-cells were further apart, and Myeloid cell types DC, monocytes, and neutrophils were close to each other.

### *scDist* detects cell-types that are different in COVID-19 patient compared to controls

We applied *scDist* to a large COVID-19 dataset (18) consisting of 1.4 million cells of 64 types from 284 PBMC samples from 196 individuals consisting of 171 COVID-19 patients and 25 healthy donors. The large number of samples of this dataset permitted further evaluation of our approach using real data rather than simulations. Specifically, we defined true distances between the two groups by computing the sum of squared log fold changes (across all genes) on the entire dataset and then estimated the distance on random samples of five cases versus five controls. Because *Augur* does not directly estimate a distance we compared the ability of the two methods to recover the relative ordering of distances observed with the ground truth. We found that *scDist* recovers the rankings better than *Augur* (**Fig 4a**; (**Fig S3**)).

**Figure 4:**
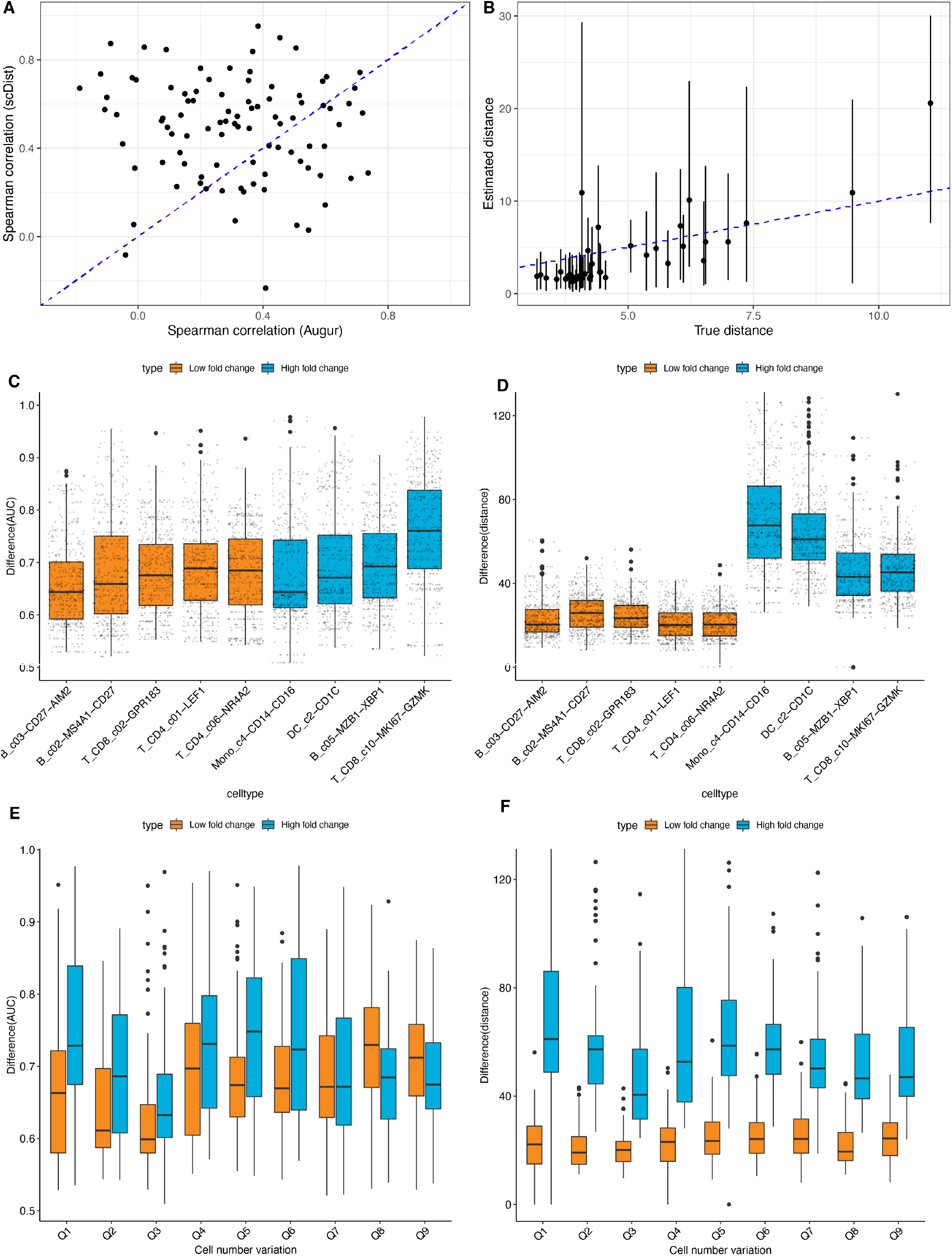
Comparison of *scDist* and *Augur* Performance Based on Real Data Simulation. **A)** Correlation between estimated ranks and true ranks for each method, with points above the diagonal line indicate better agreement of *scDist* with the true ranking. **B)** Plot of true distance vs. distances estimated with *scDist* (dashed line represents y = x). **C)** AUC values achieved by *Augur*, where color represents likely true (blue) or false (orange) positive cell types. **D)** Same as **C)**, but for distances estimated with *scDist*. **E)** AUC values achieved by *Augur* against the cell number variation in subsampled-datasets (of false positive cell types). **F)** Same as **E)**, but for distances estimated with *scDist*.

To evaluate *scDist* ‘s accuracy further, we defined a new ground truth using the entire COVID-19 dataset, consisting two groups: 4 cell types with differences between groups (true positives) and 5 cell types without differences (false positives) (**Fig S4**, Methods). We generated 1000 random samples with only five individuals per cell-type and estimated group differences using both *Augur* and *scDist. Augur* failed to accurately separate the two groups (**Fig 4c**); median difference estimates of all true positive cell types, except MK167+ CD8+T, were lower than median estimates of all true negative cell types (**Fig 4c**). In contrast, *scDist* showed a separation between *scDist* estimates between the two groups (**Fig 4d**).

Single-cell data can also exhibit dramatic sample-specific variations in the number of cells of specific cell types. This imbalance can arise from differences in collection strategies, biospecimen quality, and technical effects, and can impact the reliability of methods that do not account for sample-to-sample or individual-to-individual variations. We measured the variation in cell numbers within samples by calculating the ratio of the largest sample’s cell count to the total cell counts across all samples (Methods). *Augur* ‘s predictions were negatively impacted by this cell number variation (**Fig 4e**), indicating its increased susceptibility to false positives when sample-specific cell number variation was present (**Fig 1c**). In contrast, scDist’s estimates were robust to sample-specific cell number variation in single-cell data (**Fig 4f**).

To further demonstrate the advantage of statistical inference in the presence of individual-to-individual variation, we analyzed the smaller COVID-19 dataset (1) with only 13 samples. The original study (1) discovered differences between cases and controls in CD14+ monocytes through extensive manual inspection. *scDist* identified this same group as the most significantly perturbed cell type. *scDist* also identified two cell-types not identified in the original study, dendritic cells (DCs) and plasmacytoid dendritic cells (pDCs) (**Fig S5a**). We note that DCs induce anti-viral innate and adaptive response through antigen presentation (19). Our finding was consistent with studies reporting that DCs and pDCs are perturbed by COVID-19 infection (20; 21). In contrast, *Augur* identified red blood cells, not CD14+ monocytes, as the most perturbed cell type (**Fig S6**). Omitting the patient with the most red blood cells dropped the perturbation between infected and control cases estimated by *Augur* for red blood cells markedly (**Fig S6**), further suggesting that *Augur* predictions are clouded by patient-level variability.

### *scDist* enables the identification of genes underlying cell-specific across-condition differences

To identify transcriptomic alteration, *scDist* assigns an importance score to each gene based on its contribution to the overall perturbation (Methods). We assessed this importance score for CD14+ monocytes in small COVID-19 datasets. In this cell type, *scDist* assigned the highest importance score to genes S100A8 and S100A9 (**Fig S5b**). These genes are canonical markers of inflammation (22) that are upregulated during cytokine storm. Since patients with severe COVID-19 infections often experience cytokine storms, the result suggests that S100A8/A9 upregulation in CD14+ monocyte could be a marker of the cytokine storm (23). These two genes were reported to be upregulated in COVID-19 patients in the study of 284 samples (18).

### *scDist* identifies transcriptomic alterations associated with immunotherapy response

To demonstrate the real-world impact of *scDist*, we applied it to four published dataset used to understand patient responses to cancer immunotherapy in head and neck, bladder, and skin cancer patients respectively (24; 2; 25; 26). We found that each individual dataset was underpowered to detect differences between responders and non-responders (**Fig S7**). To potentially increase power, we combined the data from all cohorts (**Fig 5a**). However, we found that analyzing the combined data without accounting for cohortspecific variations lead to false positives. For example, responder-nonresponder differences estimated by *Augur* were highly correlated between pre-and post-treatments (**Fig 5b**), suggesting a confounding effect of cohort-specific variations. Furthermore, *Augur* predicted that most cell types were altered in both pretreatment and post-treatment samples (AUC *>* 0.5 for 41 in pre-treatment and 44 in post-treatment out of a total of 49 cell types), which is potentially due to confounding effect of cohort-specific variations.

**Figure 5:**
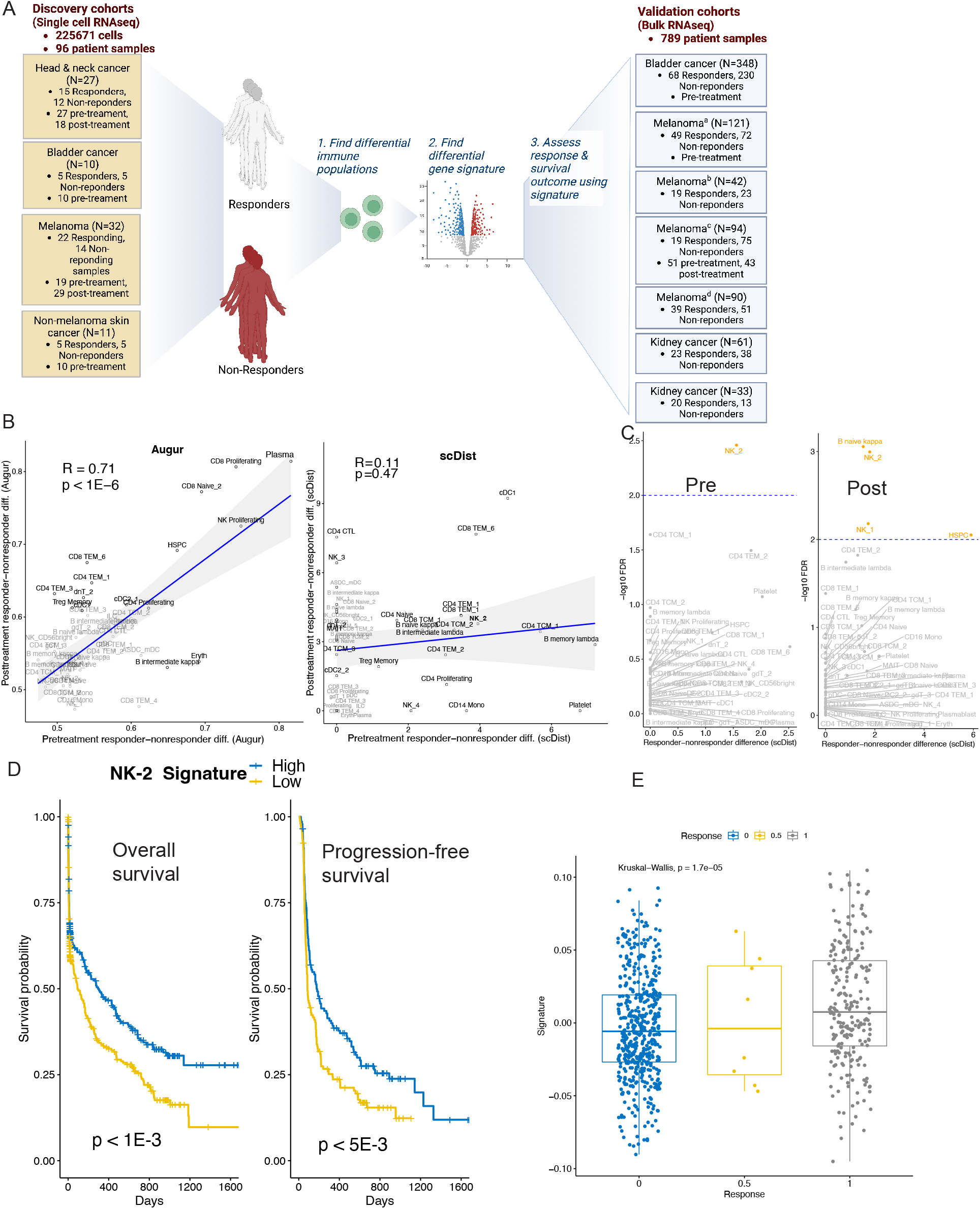
Immunotherapy Cohorts Analysis using *scDist*. **A)** Study design: discovery cohorts of four scRNA cohorts identify cell-type-specific differences and a differential gene signature between responders and non-responders. This signature was evaluated in validation cohorts of six bulk RNA-seq cohorts. **B)** Pretreatment and posttreatment sample differences were estimated using *Augur* and *scDist*. **C)** Significance of the estimated differences (*scDist*). **D)** Kaplan-Meier plots display the survival differences in anti-PD-1 therapy patients, categorized by low-risk and high-risk groups using the median value of the NK-2 signature value; overall and progression-free survival is shown. **E)** NK-2 signature levels in non-responders, partial-responders, and responders (Bulk RNA-seq cohorts).10

To account for cohort-specific variations, we ran *scDist* including an explanatory variable to the model (1) to account for cohort effects. With this approach, distance estimates were not correlated significantly between pre-and post-treatment. (**Fig 5b**). Instead, *scDist* predicted CD4-T and CD8-T altered pretreatment (**Fig S8a**), while NK, CD8-T, and B cells altered post-treatment (**Fig S8b**). Analysis of subtypes revealed FCER1G+NK cells (NK-2) were changed in both pre-treatment and post-treatment samples (**Fig 5c**). To validate this finding, we generated an NK-2 signature differential between responders and non-responders (**Fig S9** Methods) and evaluated these signatures in bulk-RNAseq immunotherapy cohorts, composing 789 patient samples (**Fig 5a**). We scored each of the 789 patient samples using the NK-2 differential signature (Methods). The NK-2 signature scores were significantly associated with overall and progression-free survival (**Fig 5d**) as well as radiology-based response (**Fig 5e**). We similarly evaluated the top *Augur* prediction. Differential signature from plasma, the top predicted cell type by *Augur*, did not show an association with the response or survival outcomes in 789 bulk transcriptomes (**Fig S10**, Methods).

### scDist is computationally efficient

A key strength of the linear modeling framework used by *scDist* is that it is efficient on large datasets. For instance, on the COVID-19 dataset with 13 samples (1), *scDist* completed the analysis in around 50 seconds, while *Augur* required 5 minutes. To better understand how runtime depends on the number of cells, we applied both methods to subsamples of the dataset that varied in size and observed that *scDist* was, on average, five-fold faster (**Fig S11**). *scDist* is also capable of scaling to millions of cells. On simulated data, *scDist* required approximately 10 minutes to fit a dataset with 1, 000, 000 cells (**Fig S12**). We also tested the sensitivity of *scDist* to the number of PCs used by comparing *D*_*K*_ for various values of *K*. We observed that the estimated distances stabilize as *K* increases (**Fig S13**), justifying *K* = 20 as a reasonable choice for most datasets.

## Discussion

The identification of cell types influenced by infections, treatments, or biological conditions is crucial for understanding their impact on human health and disease. We present *scDist*, a statistically rigorous and computationally fast method for detecting cell-type specific differences across multiple groups or conditions. By using a mixed-effects model, *scDist* estimates the difference between groups while quantifying the statistical uncertainty due to individual-to-individual variation and other sources of variability. We validated *scDist* through the unbiased recapitulation of known relationships between immune cells and demonstrated its effectiveness in mitigating false positives from patient-level and technical variations in both simulated and real datasets. Notably, *scDist* facilitates biological discoveries from scRNA cohorts, even when the number of individuals was limited, a common occurrence in human scRNA-seq datasets. We also pointed out how the detection of cell-type specific differences can be obscured by batch effects or other confounders and how the linear model used by our approach permits accounting for these.

Our study underscored the vitality of principled statistical methods in analyzing scRNA-seq data. Specifically, by using a model that permits a mathematical definition of difference and also accounts for statistical uncertainty, *scDist* can distinguish between cases in which we can detect differences and cases with not enough power. For example, in the context of COVID studies, a model sample size of five individuals per group proved adequate, given the expected substantial differences between COVID-infected patients and healthy controls. However, in immunotherapy cohorts with similar sample sizes, no significant differences were observed between responders and non-responders, suggesting the need for larger sample sizes in contexts where between-group differences are expected to be less pronounced. In this case, our model-based approach permitted the integration of data from multiple studies to provide enough power to permit biological insights related to transcriptomic alterations associated with immunotherapy response.

We have shown that *scDist* greatly outperforms current approaches, however, there are several ways it could be improved in future versions. First, *scDist* requires pre-specification of the number of principal components to estimate distance. In practice, we observed that the estimated distance is stable as the number of PCs varies between 20 and 50 (**Fig S13**). An adaptive approach to selecting the number of PCs could improve our approach. Second, *scDist* does not currently account for unknown batch effects. We note that current approaches for removing known and unknown batch effects in bulk RNA-seq data (14; 27), are likely to inadvertently eliminate the biological signal of interest in this specific application (**Fig S1**). Third, *scDist* relies on the accurate annotation of cell-types; therefore, inaccuracies in existing clustering and annotation approaches could impact its performance. A future version could add a way to identify perturbed regions of gene expression space without relying on a predefined clustering (4; 5; 6). Fourth, although Pearson residual-based normalized counts (15; 16) is recommended input for *scDist*, if the data available was normalized by another, sub-optimal, approach, *scDist* ‘s performances could be affected. A future version could adapt the model and estimation procedure so that *scDist* can be directly applied to the counts, and avoid potential problems introduced by normalization.

We believe that *scDist* will have extensive utility, as the comparison of single-cell experiments between groups is a common task across a range of research and clinical applications. In this study, we have focused on examining discrete phenotypes, such as infected versus non-infected (in COVID-19 studies) and responders vs. non-responders to checkpoint inhibitors. However, the versatility of our framework allows for extension to experiments involving continuous phenotypes or conditions, such as height, survival, and exposure levels, to name a few. As single-cell datasets continue to grow in size and complexity, *scDist* will enable rigorous and reliable insights into cellular perturbations with implications to human health and disease.

## Acknowledgements

PBN is supported by NIH T32CA009337. ADS received support from K99CA248953, the Michelson Foundation, and was partially supported by the UNM Comprehensive Cancer Center Support Grant NCI P30CA118100. Figures 2 and 5 were created using BioRender: Biorender.com. We express our gratitude to Adrienne M. Luoma, Shengbao Suo, and Kai W. Wucherpfennig for providing the scRNA data (24). We also thank Zexian Zeng for assistance with downloading and accessing the bulk RNA-seq dataset.

## Methods

### Normalization

Our method takes as input a normalized count matrix (with corresponding cell type annotations). We recommend using *scTransform* (16) to normalize, although the method is compatible with any normalization approach. Let *y*_*ijg*_ be the UMI counts for gene 1 ≥ *g* ≥ *G* in cell *i* from sample *j. scTransform* fits the following model:

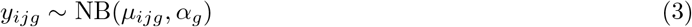

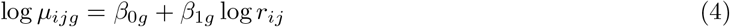

where *r*_*ij*_ is the total number of UMI counts for the particular cell. The normalized counts are given by the Pearson residuals of the above model:

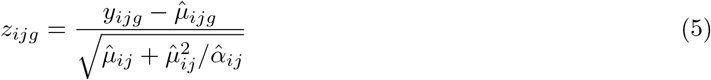

### Distance in normalized expression space

In this section, we describe the inferential procedure of *scDist* for cases without additional covariates. However, the procedure can be generalized to the full model with arbitrary covariates (design matrix) incorporating random and fixed effects, as well as nested-effect mixed models. For a given cell type, we model the *G*-dimensional vector of normalized counts as

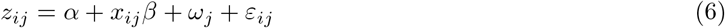

where *α, β* ∈ ℝ^*G*^, *x*_*ij*_ is a binary indicator of condition, *ω*_*j*_ ∼ 𝒩 (0, *τ* ^2^*I*_*G*_), and *ε*_*ij*_ ∼ 𝒩 (0, *σ*^2^*I*_*G*_). The quantity of interest is the Euclidean distance between condition means *α* and *α* + *β*:

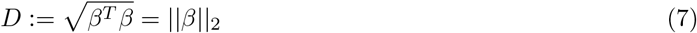

If *U* ∈ ℝ ^*G×G*^ is an orthonormal matrix, we can apply *U* to equation (6) to obtain the *transformed model* :

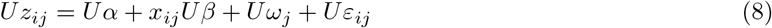

Since *U* is orthogonal, *Uω*_*j*_ and *Uε*_*ij*_ still have spherical normal distributions. We also have that

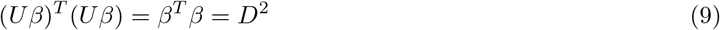

This means that the distance in the transformed model is the same as in the original model. As mentioned earlier, our goal is to find *U* such that

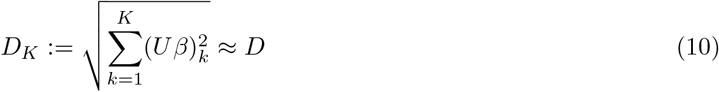

with *K* ≪ *G*.

Let *Z* ∈ ℝ^*n×G*^ be the matrix with rows *z*_*ij*_ (where *n* is the total number of cells). Intuitively, we want to choose a *U* such that the projection of *z*_*ij*_ onto the first *K* rows of *U* (*u*_1_, …, *u*_*K*_ ∈ ℝ^*G*^) minimizes the reconstruction error

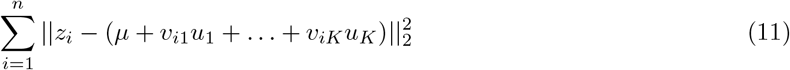

where *µ* ∈ ℝ^*G*^ is a shift vector and (*v*_*ik*_) ∈ ℝ^*n×K*^ is a matrix of coefficients. It can be shown that the PCA of *Z* yields the (orthornormal) *u*_1_, …, *u*_*K*_ that minimizes this reconstruction error (28).

### Inference

Given an estimator 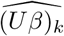 of (*Uβ*)_*k*_, a naive estimator of *D*_*K*_ is given by taking the square root of the sum of squared estimates:

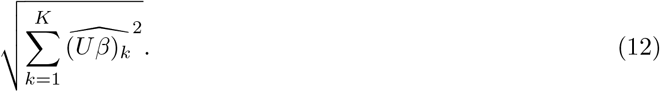

However, this estimator can have significant upward bias due to sampling variability. For instance, even if the true distance is 0, 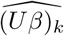 is unlikely to be exactly zero, and that noise becomes strictly positive when squaring.

To account for this, we apply a post-hoc Bayesian procedure to the 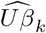 to shrink them towards zero before computing the sum of squares. In particular, we adopt the spike slab model of (17)

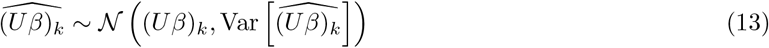

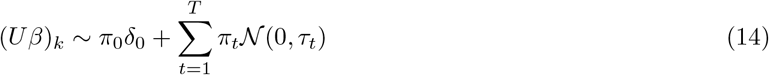

where 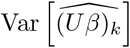 is the variance of the estimator 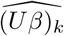 is a point mass at 0, and *π*_0_, *π*_1_, … *π*_*T*_ are mixing weights (that is, they are non-negative and sum to 1). (17) provides a fast empirical Bayes approach to estimate the mixing weights and obtain posterior samples of (*Uβ*)_*k*_. Then samples from the posterior of *D*_*K*_ are obtained by applying the formula (12) to the posterior samples of (*Uβ*)_*k*_. We then summarize the posterior distribution by reporting the median and other quantiles. Advantages of this particular specification is that the amount of shrinkage depends on the uncertainty in the initial estimate of (*Uβ*)_*k*_.

We use the following procedure to obtain 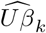:

1. Use the matrix of PCA loadings as a plug in estimator for *U*. Then *Uz*_*ij*_ is the vector of PC scores for cell *i* in sample *j*.
2. Estimate (*Uβ*)_*k*_ by using *lme4* (29) to fit the model (6) using the PC scores corresponding to the *k*-th loading (i.e., each dimension is fit independently).

Note that only the first *K* rows of *U* need to be stored.

We are particularly interested in testing the null hypothesis of *D*_*K*_ = 0 against the alternative *D*_*d*_ > 0. Because the null hypothesis corresponds to (*Uβ*)_*k*_ = 0 for all 1 ≥ *k* ≥ *d*, we can use the sum of individual Wald statistics as our test statistic:

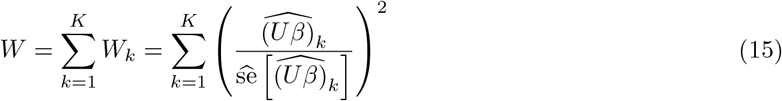

Under the null hypothesis that (*Uβ*)_*k*_ = 0, *W*_*k*_ can be approximated by a 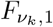 distribution. *ν*_*k*_ is estimated using Satterthwaite’s approximation in *lmerTest*. This implies that

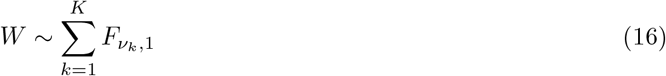

under the null. Moreover, the *W*_*k*_ are independent because we have assumed that covariance matrices for the sample and cell-level noise are multiples of the identity. Equation (16) is not a known distribution but quantiles can be approximated using Monte Carlo samples. To make this precise, let *W*_1_, …, *W*_*M*_ be draws from equation (16), where *M* = 10^5^ and let *W* ^*∗*^ be the value of equation (15) (i.e., the actual test statistic). Then the empirical *p*-value (30) is computed as

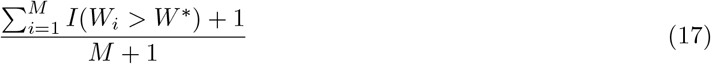

### Choosing the number of principal components

An important choice in *scDist* is the number of principal components *d*. If *d* is chosen too small, then estimation accuracy may suffer as the first few PCs may not capture enough of the distance. On the other hand, if *d* is chosen too large then the power may suffer as a majority of the PCs will simply be capturing random noise (and adding to degrees of freedom to the Wald statistic). Moreover, it is important that *d* is chosen a priori, as choosing the *d* that produces the lowest *p*-values is akin to *p*-hacking.

If the model is correctly specified then it is reasonable to choose *d* = *J* − 1, where *J* is the number of samples (or patients). To see why, notice that the mean expression in sample 1 ≥ *j* ≥ *J* is

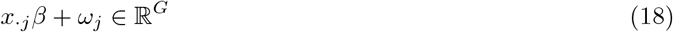

In particular, the *J* sample means lie on a (*J* − 1)-dimensional subspace in ℝ^*G*^. Under the assumption that the condition difference and sample-level variability is larger than the error variance *σ*^2^, we should expect that the first *J −* 1 PC vectors capture all of the variance due to differences in sample means.

In practice, however, the model can not be expected to be correctly specified. For this reason, we find that *d* = 20 is a reasonable choice when the number of samples is small (as is usually the case in scRNA-seq) and *d* = 50 for datasets with a large number of samples. This is line with other single-cell methods, where the number of PCs retained is usually between 20 and 50.

### Feature importance

To better understand the genes that drive the observed difference in the CD14+ monocytes we define a *gene importance score*. For 1 ≥ *k* ≥ *d* and 1 ≥ *g* ≥ *G*, the *k*-th importance score for gene *g* is |*U*_*kg*_| *β*_*g*_. In other words, the importance score is the absolute value of the gene’s *k*-th PC loading times its expression difference between the two conditions.

### Simulated single-cell data

We test the method on data generated from model equation (6). To ensure that the “true” distance is *D*, we use the ℝ package *uniformly* (31) to draw *β* from the surface of the sphere of radius *D* in ℝ^*G*^. The data in **Fig 1c** and **Fig 3c** are obtained by setting *β* = 0 and *σ*^2^ = 1 and varying *τ* ^2^ between 0 and 1.

### Semi-simulated COVID-19 data

COVID-19 patient data for the analysis was obtained from (18), containing 1.4 million cells of 64 types from 284 PBMC samples collected from 196 individuals, including 171 COVID-19 patients and 25 healthy donors.

#### Ground truth

To derive a reliable ground truth, we compared the 171 COVID-19 patients with the 25 healthy controls. To ensure the robustness of ground truth, we randomly divided the entire dataset into two non-overlapping groups and assessed the impact of patient-specific variation on the robustness of the ground truth perturbation estimates. We calculated log fold changes for each gene (between case and control) by taking the difference of condition means on the log(1 + *x*) transformed data and defined the ground truth distance as the sum of squared log fold changes across all genes. We compared the ground truth distances between the two groups. The high correlation of the results (*R* = 0.7, *P <* 2.2 *×*10^*−* 16^) suggests that the effects of patient-specific variation on perturbations were largely mitigated by the large sample sizes of the dataset, thus making the fold change estimates robust to patient-specific variation. Among the cell types with a sufficient number of cells, we selected four cell types with high fold changes as our true positive cell types and five cell types (with a large number of cells) with low fold changes as false positive cell types.

Using the ground truth we performed two separate simulation analyses:

1. <u>*Simulation analyses I (Fig 4a,b)*</u>: Using one half of the dataset (712621 cells, 132 case samples, 20 control samples), we created 100 subsamples consisting of 5 cases and 5 controls. For each subsample, we applied both *scDist* and *Augur* to estimate perturbation/distance between cases and controls for each cell type. Then we computed the correlation between the ground truth ranking (ordering cells by sum of log fold changes on the whole dataset) and the ranking obtained by both methods. For *scDist*, we restricted to cell types that had a non-zero distance estimate in each subsample and for *Augur* we restricted to cell types that had an AUC greater than 0.5 (**Fig 4a**). For **Fig 4b**, we took the mean estimated distance across subsamples for which the given cell type had a non-zero distance estimate. This is because in some subsamples a given cell type could be completely absent.
2. <u>*Simulation analyses II (Fig 4c-f)*</u>: We subsampled the COVID-19 cohort with 284 samples (284 PBMC samples from 196 individuals: 171 with COVID-19 infection and 25 healthy controls) to create 1,000 downsampled cohorts, each containing samples from 10 individuals (5 with COVID-19 and 5 healthy controls). We randomly selected each sample from the downsampled cohort, further downsampled the number of cells for each cell type, and selected them from the original COVID-19 cohort. This downsampling procedure increases both cohort variability and cell-number variations.

#### Performance Evaluation in Subsampled Cohorts

We applied *scDist* and *Augur* to each subsampled cohort, comparing the results for true positive and false positive cell types. We partitioned the sampled cohorts into 10 groups based on cell-number variation, defined as the number of cells in a sample with the highest number of cells for false-negative cell types divided by the average number of cells in cell types. This procedure highlights the vulnerability of computational methods to cell number variation, particularly in negative cell types.

### Analysis of immunotherapy cohorts

#### Data collection

We obtained single-cell data from four cohorts (24; 2; 25; 26), including expression counts and patient response information.

#### Pre-processing

To ensure uniform processing and annotation across the four scRNA cohorts, we analyzed CD45+ cells (removing CD45-cells) in each cohort and annotated cells using Azimuth (32) with reference provided for CD45+ cells.

#### Model to account for cohort and sample variance

To account for cohort-specific and sample-specific batch effects, *scDist* modeled the normalized gene expression as:

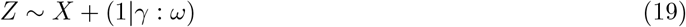

Here, Z represents the normalized count matrix, X denotes the binary indicator of condition (responder =1, non-responder=0); *γ* and *ω* are cohort and sample level random effects, and (1 *γ* : *ω*) models nested effects of samples within cohorts. The inference procedure for distance, its variance, and significance for the model with multiple cohorts is analogous to the single-cohort model.

#### Signature

We estimated the signature in the *NK-2* cell type using differential expression between responders and non-responders. To account for cohort-specific and patient-specific effects in differential expression estimation, we employed a linear mixed model described above for estimating distances, performing inference for each gene separately. The coefficient of *X* inferred from the linear mixed models was used as the estimate of differential expression:

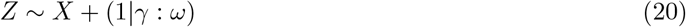

Here, Z represents the normalized count matrix, X denotes the binary indicator of condition (responder =1, non-responder=0); *γ* and *ω* are cohort and sample level random effects, and (1 *γ* : *ω*) models nested effects of samples within cohorts.

#### Bulk RNA-seq cohorts

We obtained bulk RNA-seq data from seven cancer cohorts (33; 34; 35; 36; 37; 38; 39), comprising a total of 789 patients. Within each cohort, we converted counts of each gene to TPM and normalized them to zero mean and unit standard deviation. We collected survival outcomes (both progression-free and overall) and radiologic-based responses (partial/complete responders and non-responders with stable/progressive disease) for each patient.

#### Evaluation of signature in bulk RNA-seq cohorts

We scored each bulk transcriptome (sample) for the signature using the strategy described in (40). Specifically, the score was defined as the Spearman correlation between the normalized expression and differential expression in the signature. We stratified patients into two groups using the median score for patient stratification. Kaplan-Meier plots were generated using these stratifications, and the significance of survival differences was assessed using the log-rank test. To demonstrate the association of signature levels with radiological response, we plotted signature levels separately for non-responders, partial-responders, and responders.

#### Evaluating *Augur* Signature in Bulk RNA-Seq Cohorts

A differential signature was derived for *Augur* ‘s top prediction, plasma cells, using a procedure analogous to the one described above for *scDist*. This plasma signature was then assessed in bulk RNA-seq cohorts following the same evaluation strategy as applied to the *scDist* signature.

### Software availability and data

*scDist* is available as a R package and can be downloaded from GitHub: github.com/phillipnicol/scDist

The repository also includes scripts to replicate some of the figures and a demo of *scDist* using simulated data.

Table 1 gives a list of the datasets used in each figure, as well as details about how the datasets can be obtained.

## Supplementary Materials

### A Supplementary Figures

**Figure S1:**
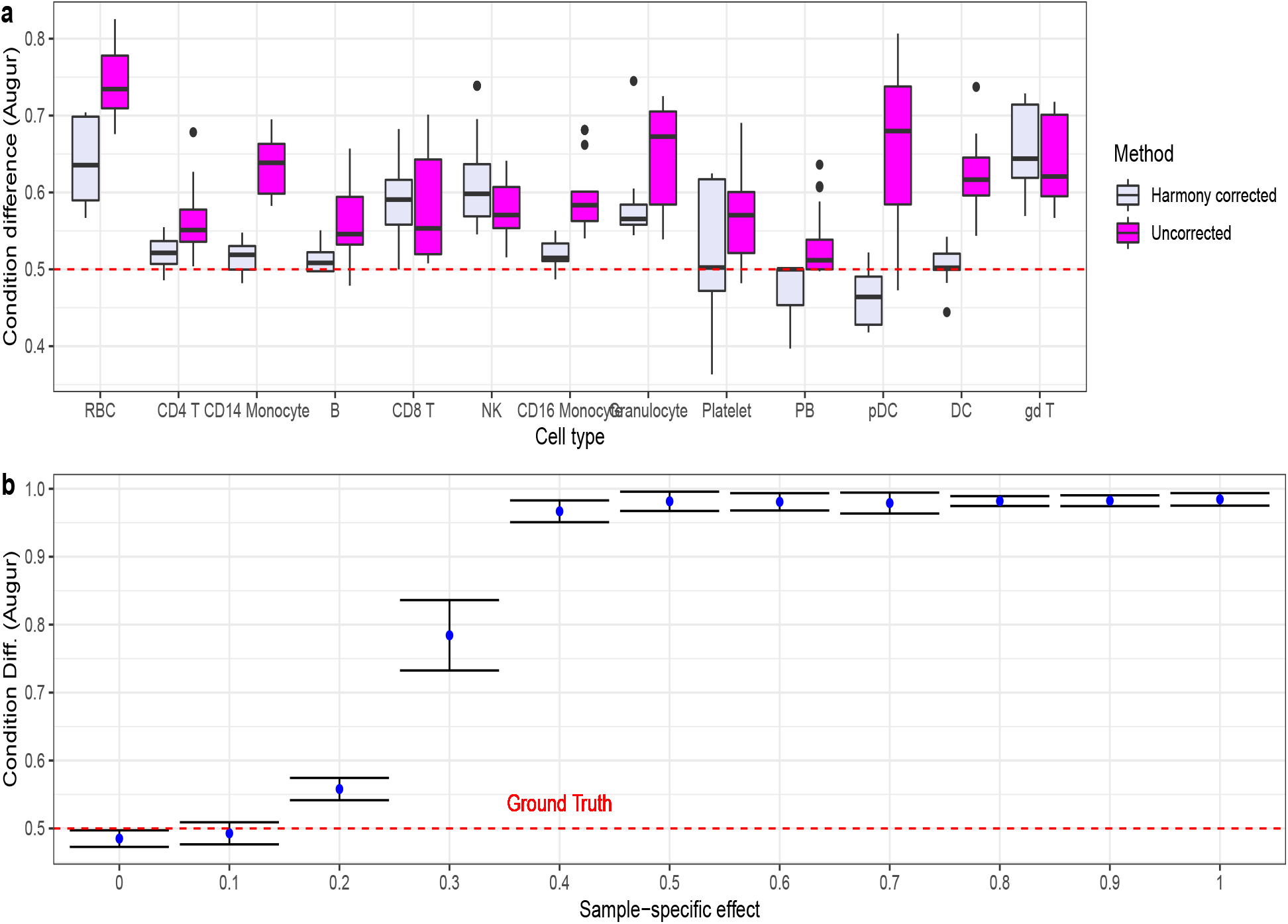
Repeating the analysis in Figure 1a and Figure 1c using batch correction as a pre-processing step. Each patient was treated as a batch and Harmony (14) was used to obtain batch-corrected PC scores. *Augur* was then applied to the PC scores.

**Figure S2:**
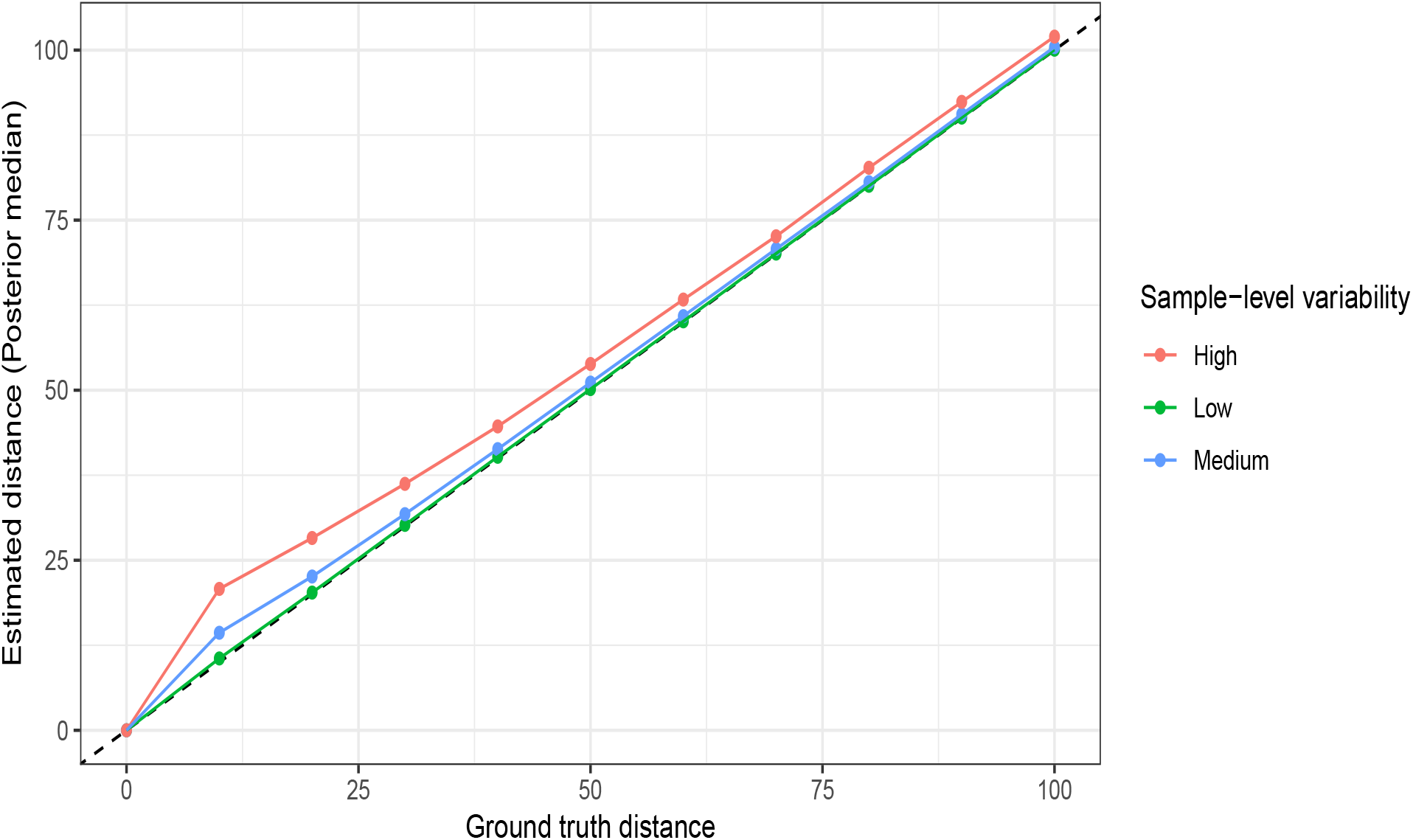
*scDist* recovers ground truth distance on simulated data. For each ground truth distance, the median estimated distance across 100 simulated distances is reported. Data was generated with *G* = 1000, 5 patients per condition, 50 cells per patient, and a patient level random effect *τ* ^2^ = 1.

**Figure S3:**
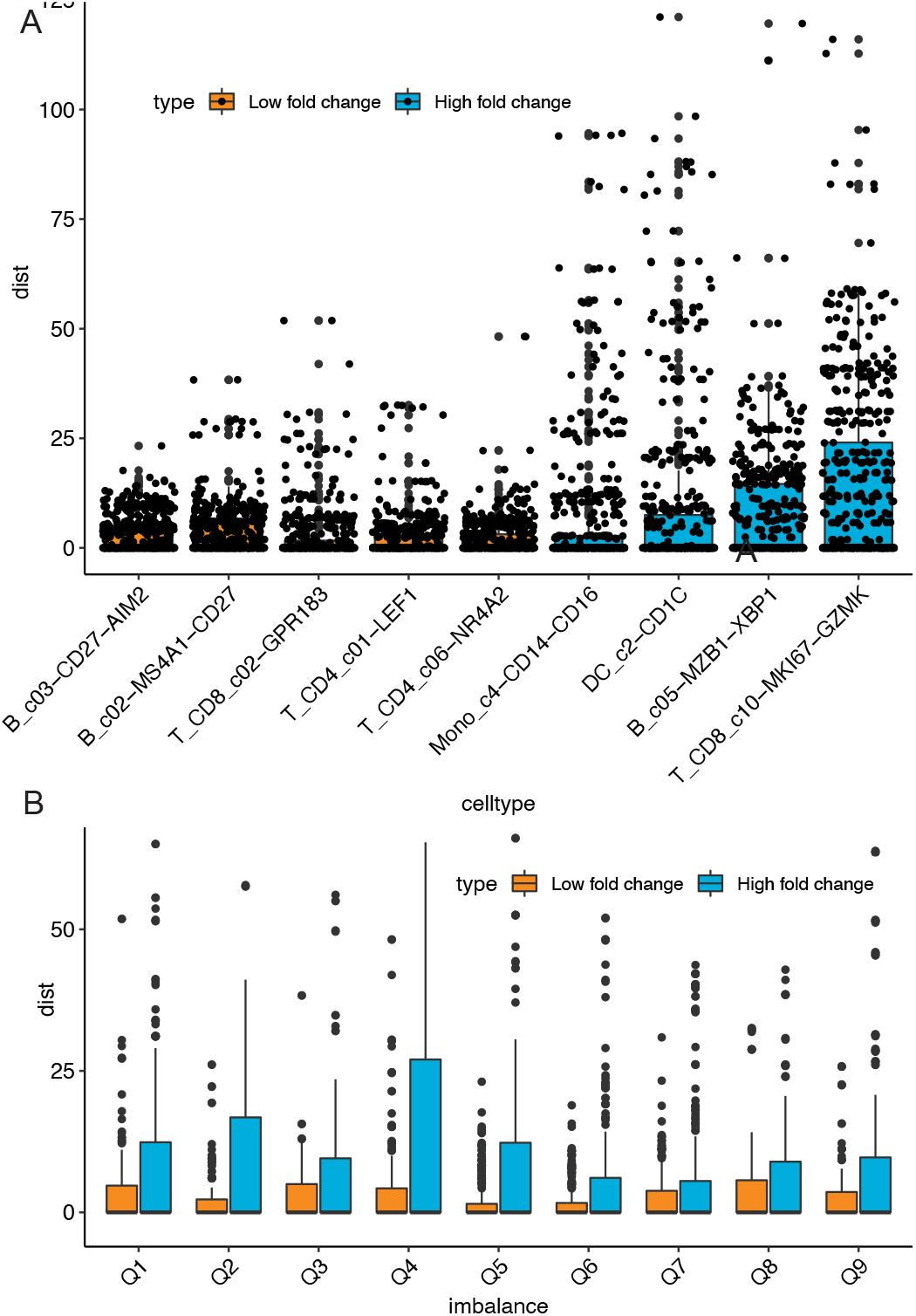
**A**. Median *scDist* distances with Bayesian correction, depicting probable true (blue) and false (orange) positive cell types. **B**. Median distances estimated by *scDist* illustrating the influence of cell number variation in subsampled datasets (of false positive cell types).

**Figure S4:**
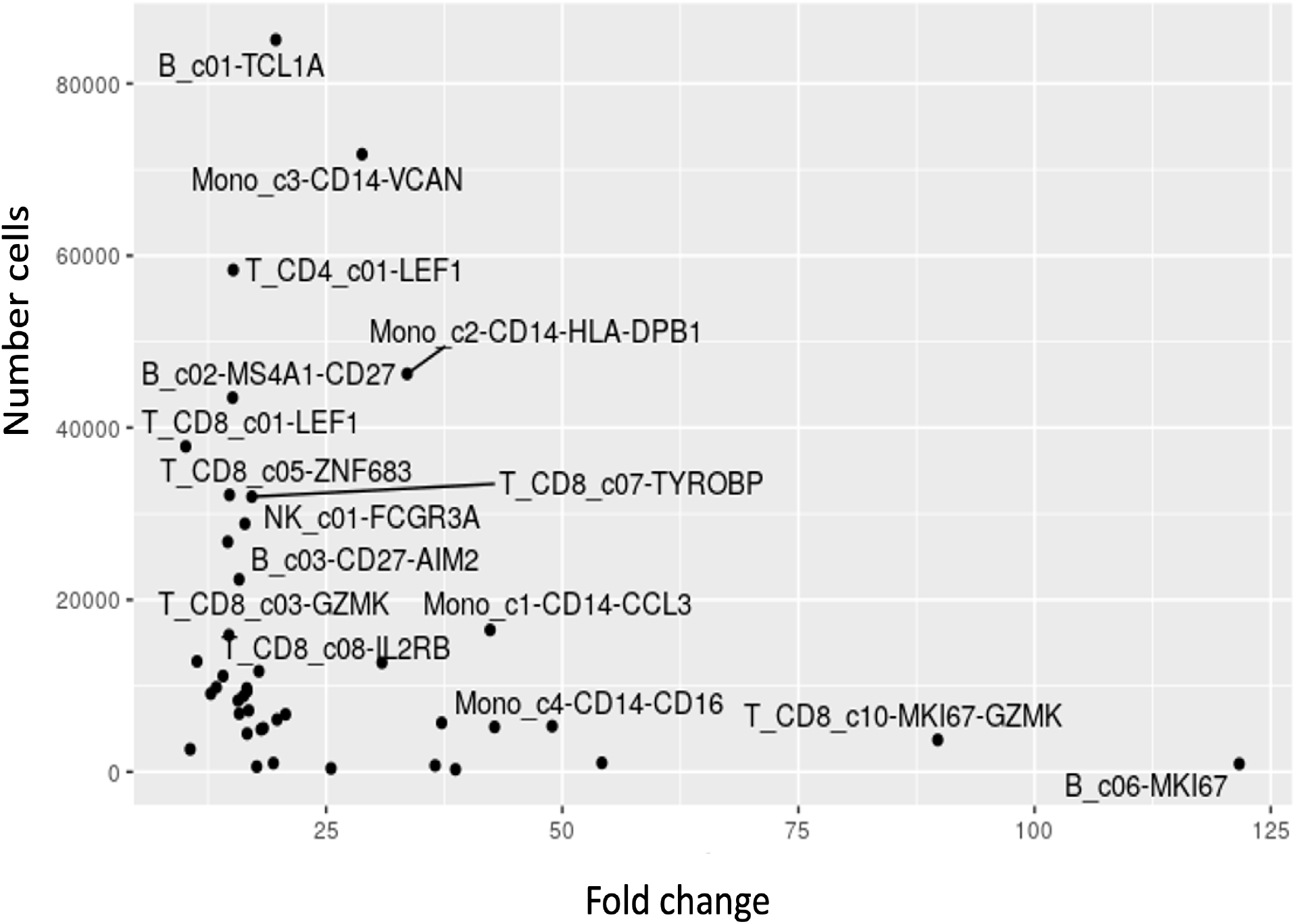
Comparative Analysis of Ren. et. al. COVID-19 Single-Cell Cohort (284 Samples). The plot displays the average gene fold changes between COVID-19 and control samples for various cell types, with the number of cells in the cohort on the y-axis.

**Figure S5:**
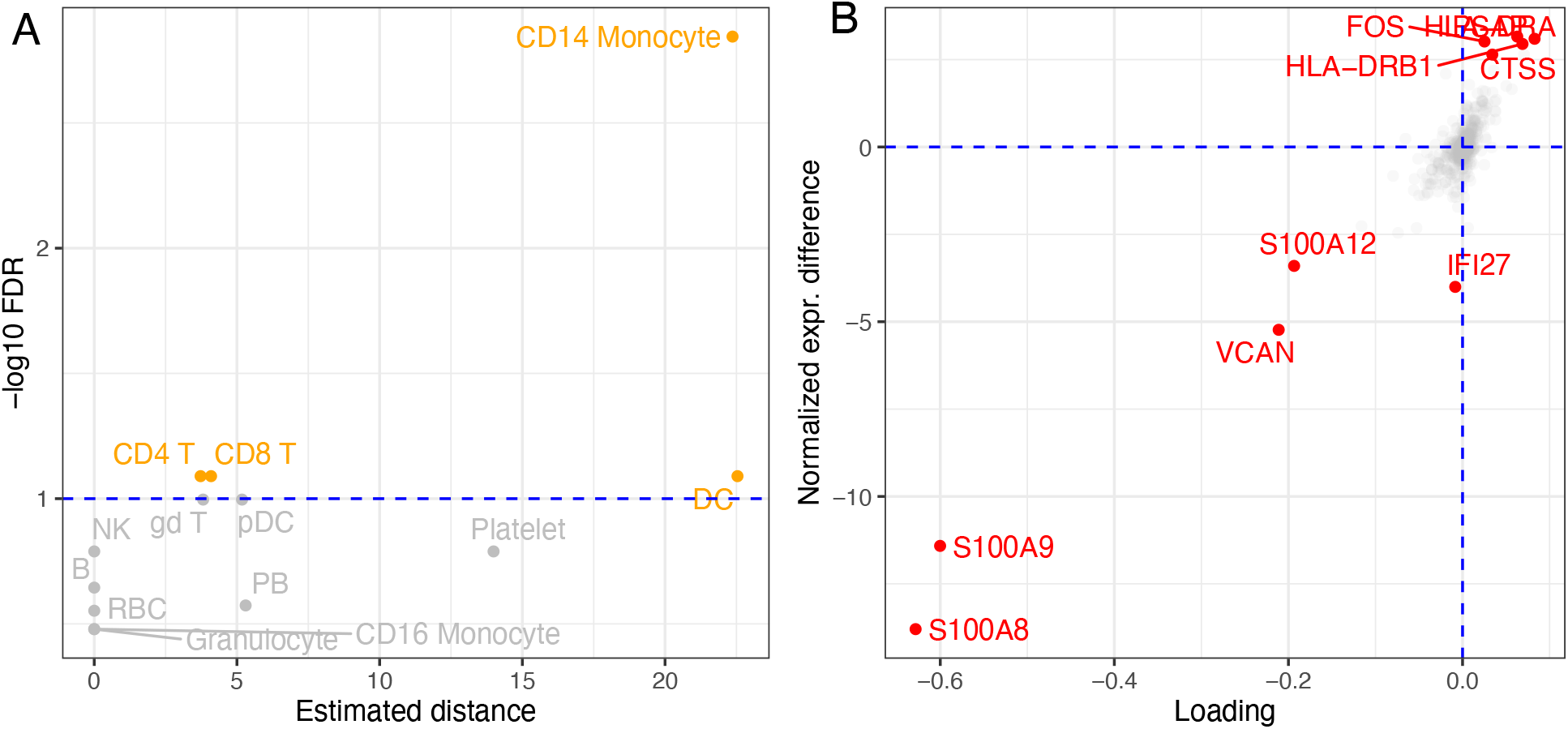
**A**. Estimated difference for each cell type in the COVID-19 data is plotted against the false discovery rate (FDR). **B**. Focusing on CD14+ monocytes, the PC1 weight for each gene is plotted against its expression difference (comparing cases and controls); red-colored genes have the highest contribution to the identified perturbation in CD14+ monocytes.

**Figure S6:**
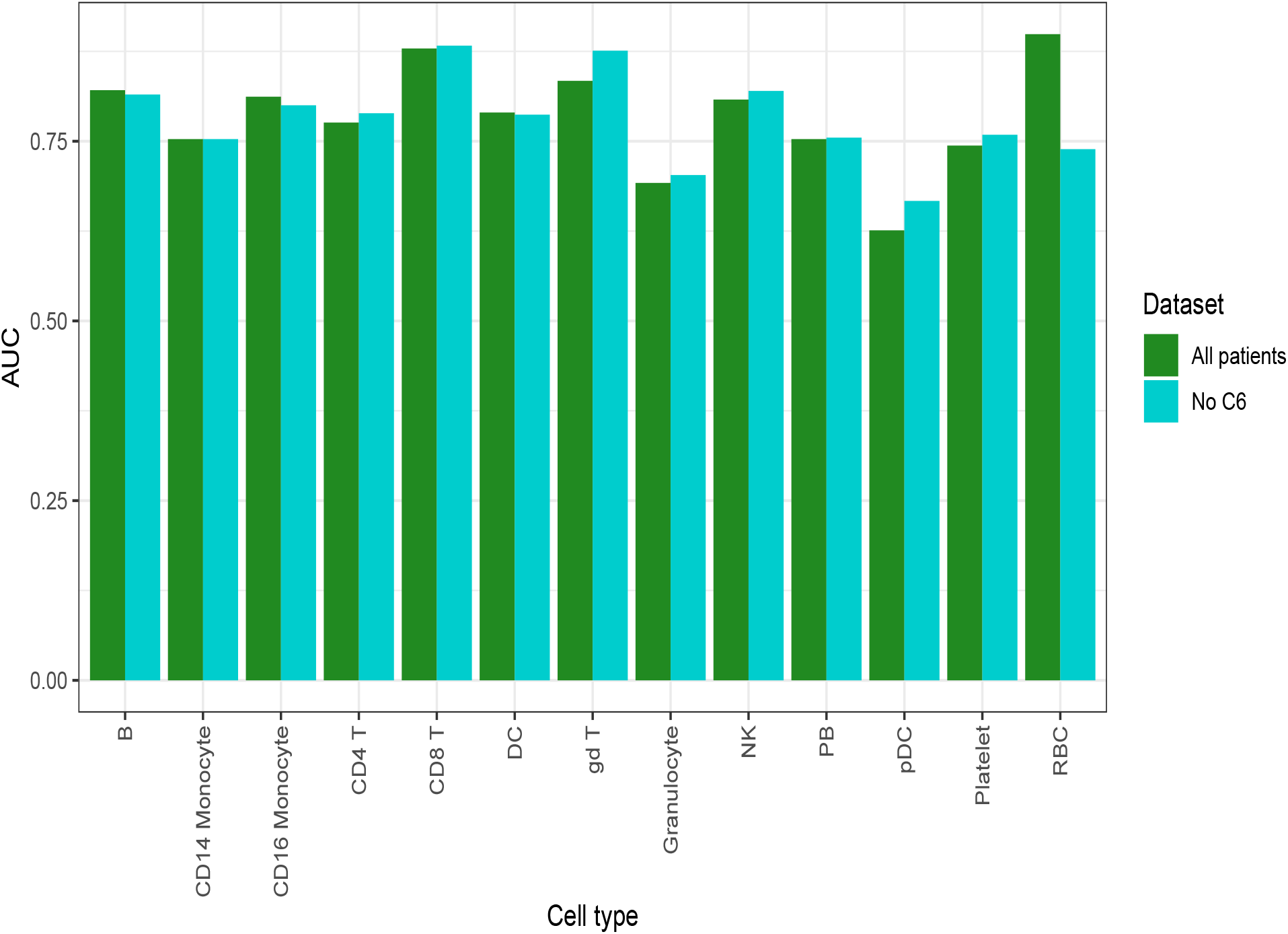
Applying *Augur* to the (1) data with the sixth case (patient C6) removed.

**Figure S7:**
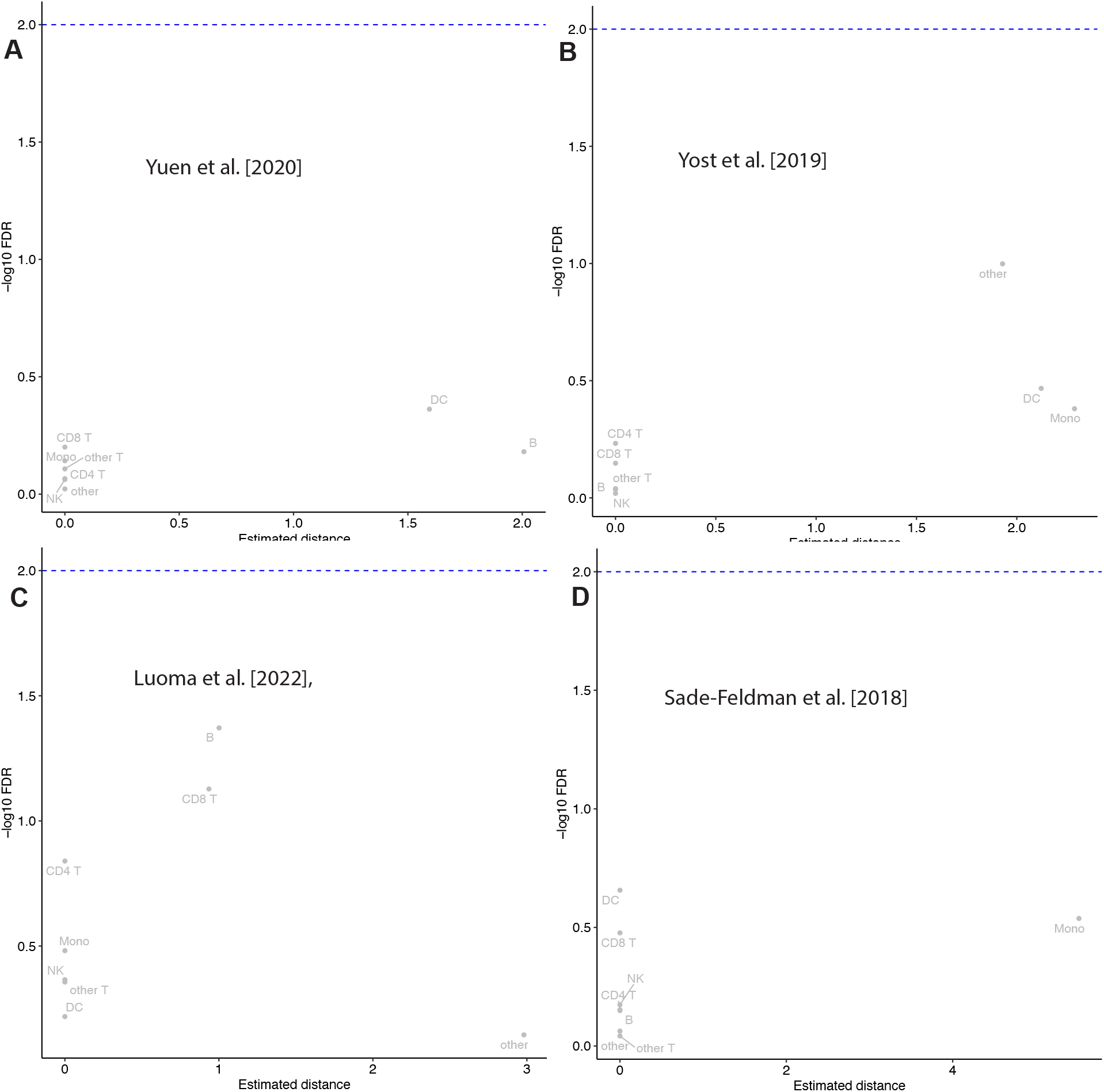
Analysis of Responder and Non-responder Groups Using *scDist* in Four Single-Cell Cohorts. Application of *scDist* to compare responders and non-responders in four independent single-cell cohorts. No significant cell types were identified in any of the cohorts.

**Figure S8:**
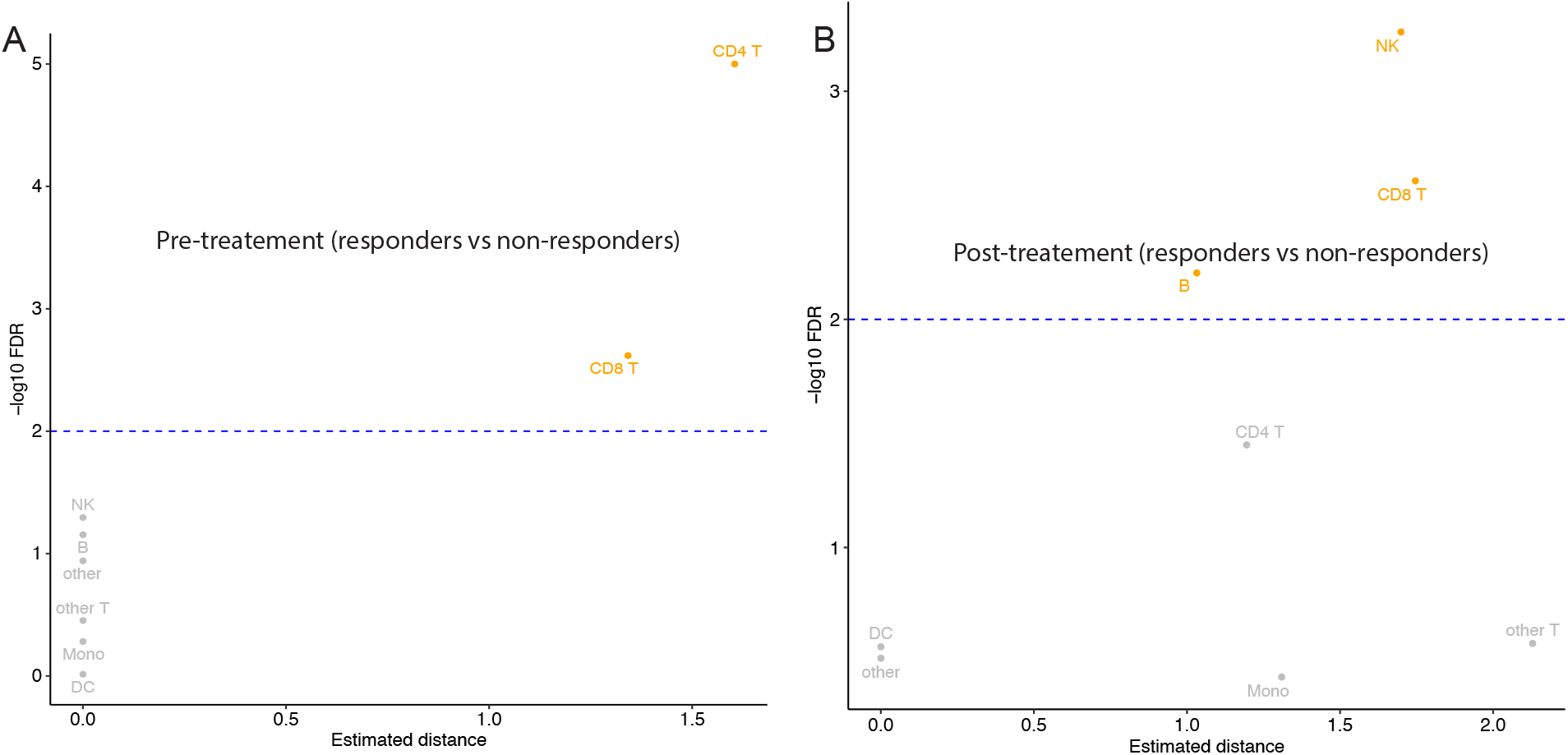
Comparing Responders and Nonresponders Using *scDist* in Integrated Single-Cell Cohorts. Application of *scDist* to assess differences between responders and non-responders in an integrated analysis of single-cell cohorts. **A)** Pretreatment differences and their significance. **B)** Posttreatment differences and their significance.

**Figure S9:**
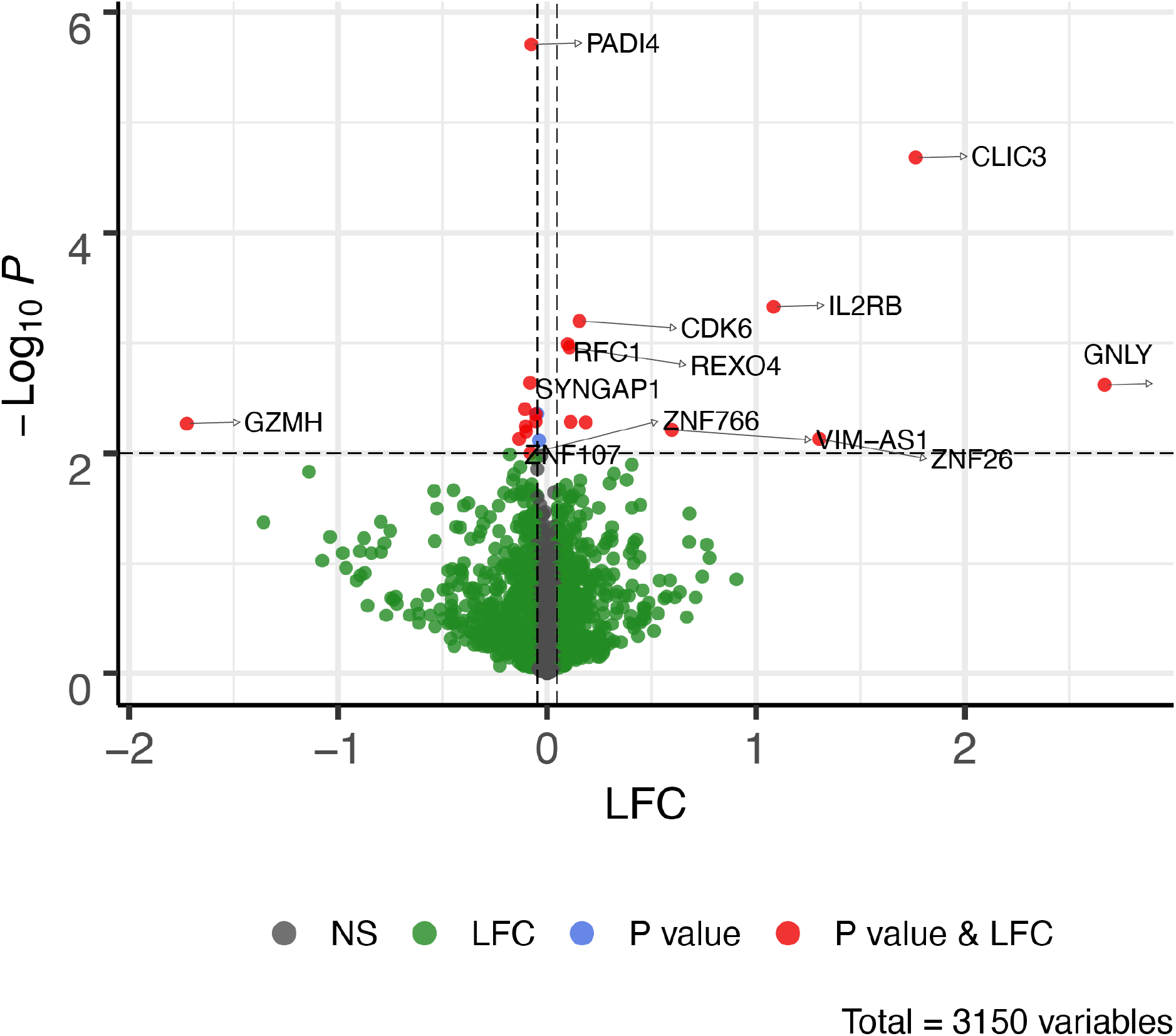
Differential expression analysis of responders and non-responders in NK-2 cells from integrated single-cell cohorts. A volcano plot illustrating the differential expression between responders and non-responders within NK-2 cells from integrated single-cell cohorts. The plot displays both the magnitude of differential expression and its statistical significance.

**Figure S10:**
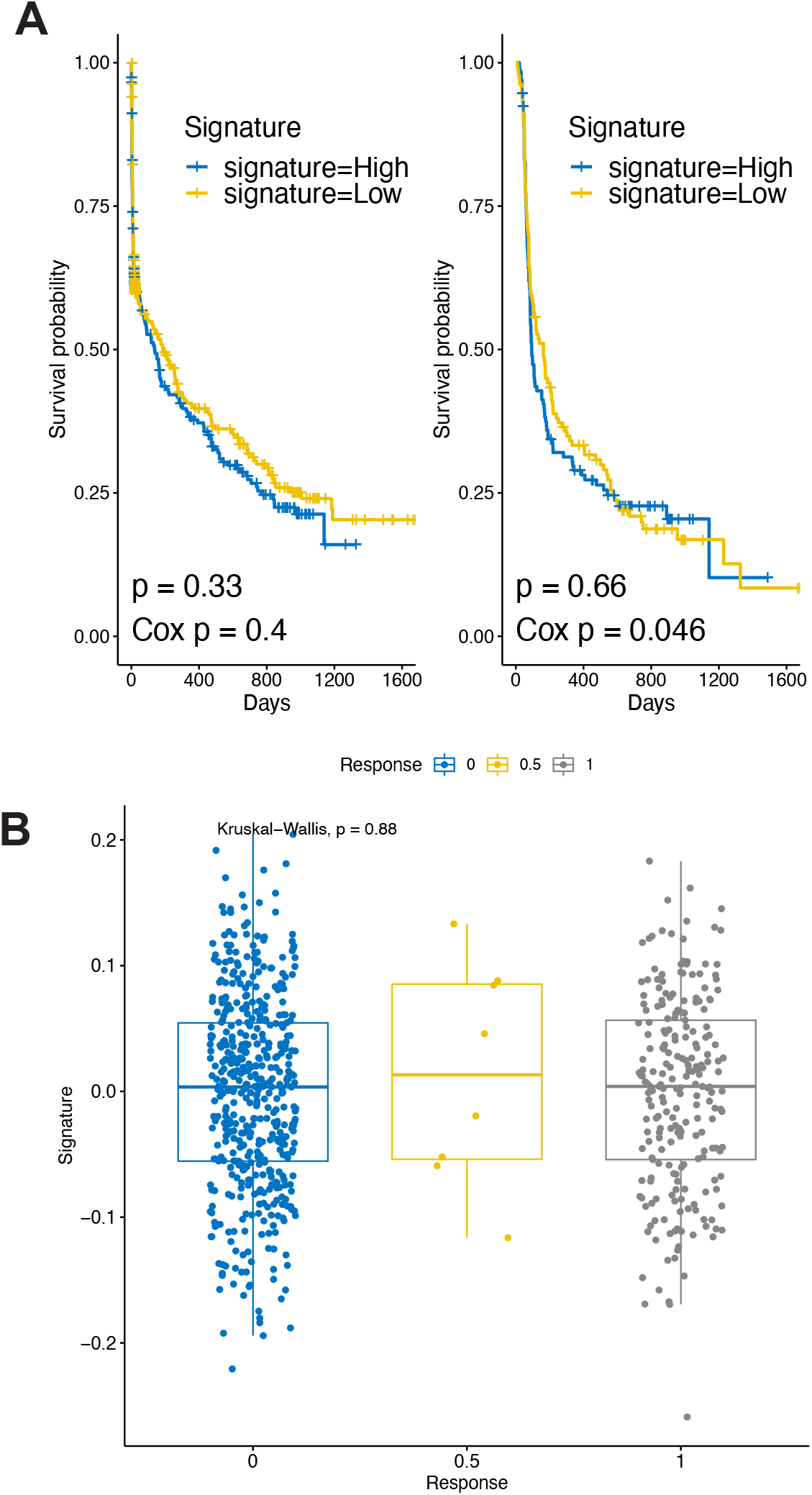
A. Kaplan-Meier plots display the survival differences in anti-PD-1 therapy patients, categorized by low-risk and high-risk groups using the median value of the Plasma signature value; overall and progression-free survival is shown. **B)** Plasma signature levels in non-responders, partial-responders, and responders (Bulk RNA-seq cohorts).

**Figure S11:**
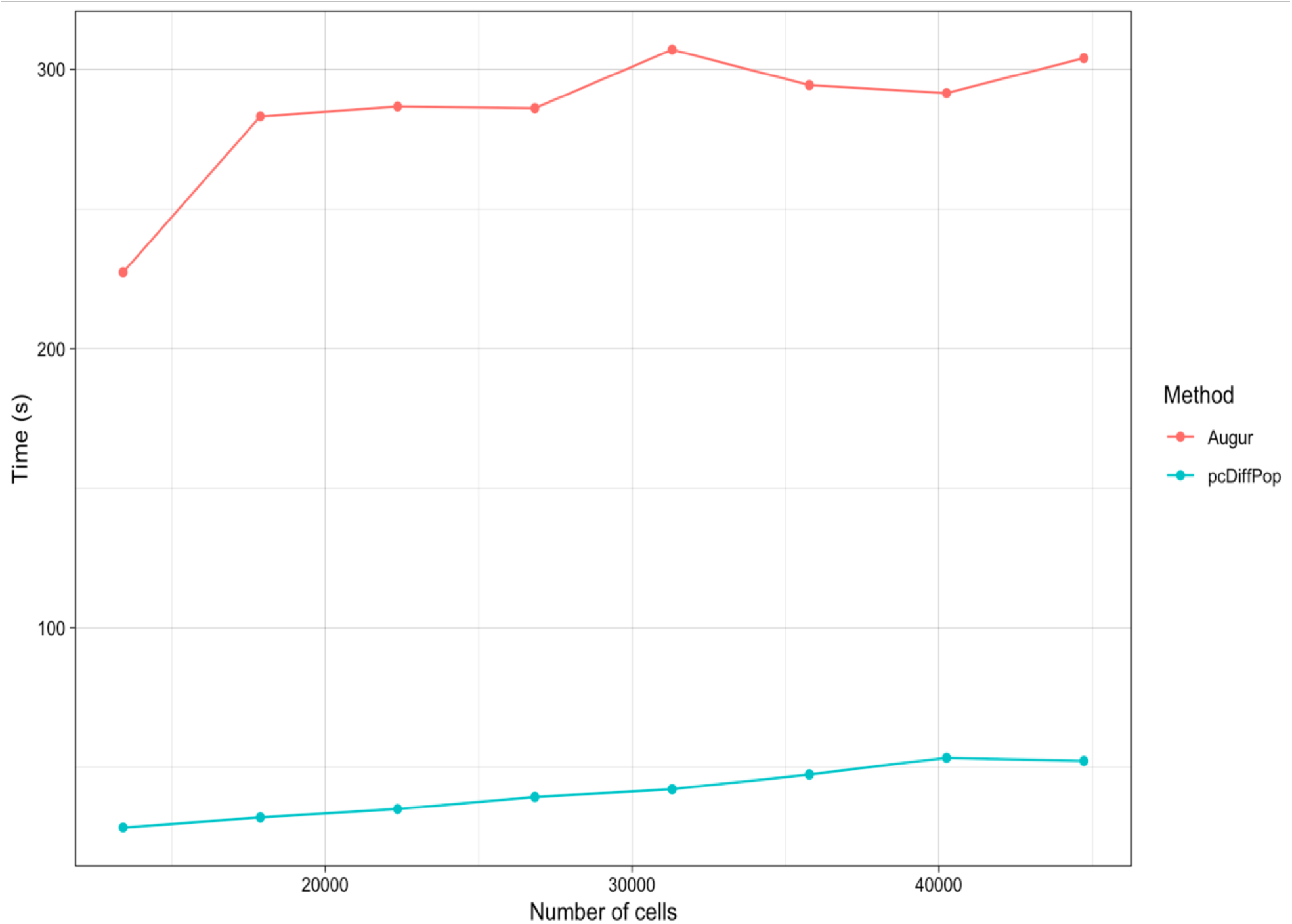
Comparison of the run-time between *Augur* and *scDist* on random subsets of the (1) data.

**Figure S12:**
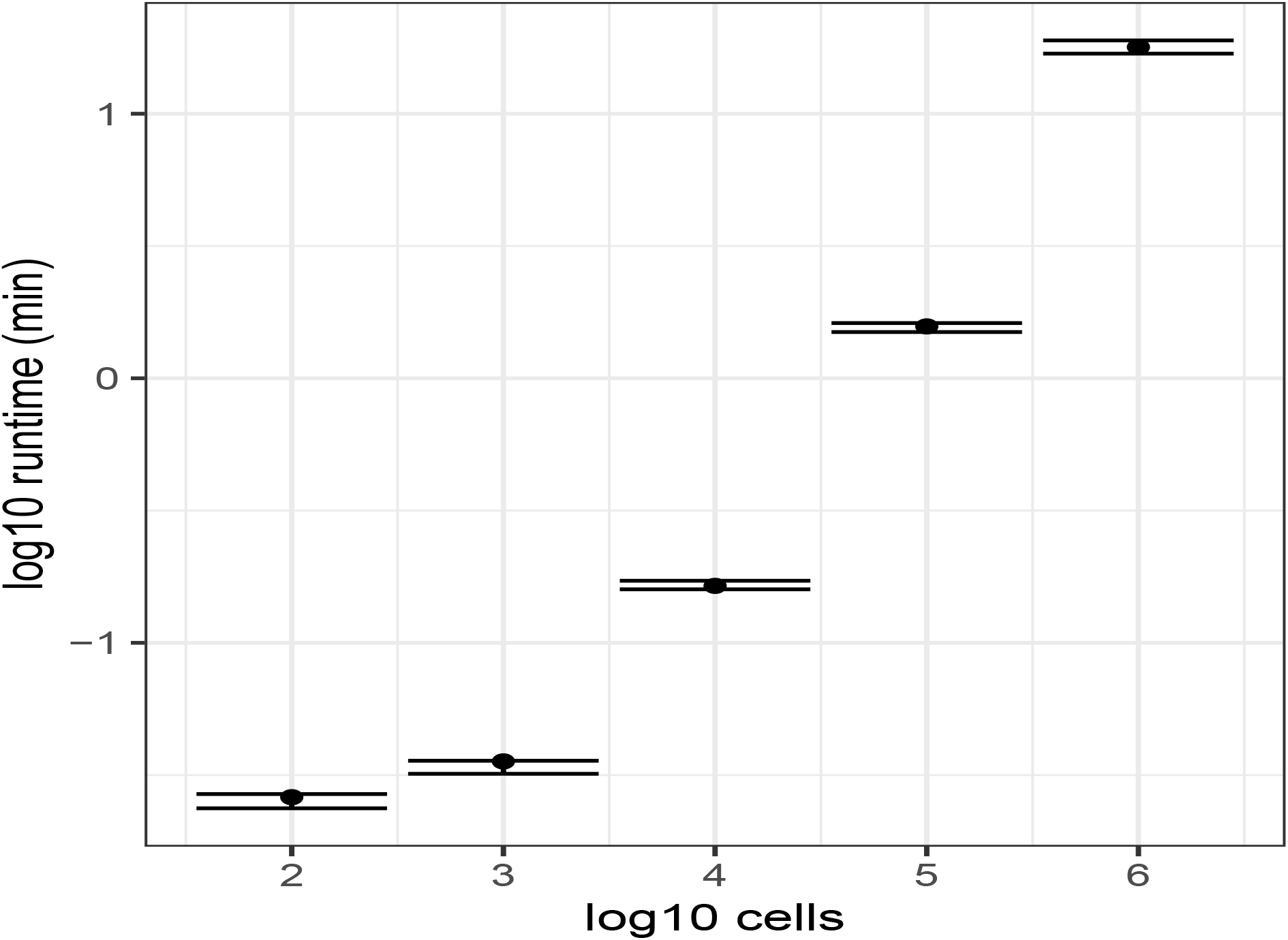
Log-log plot of runtime for *scDist* for simulated datasets with varying number of cells. Median and interquartile range across 10 iterations are reported for each size. 10 patients were simulated with an equal number of cells per patient. Data was generated with *G* = 1000, 5 patients per condition, 50 cells per patient, and a patient level random effect *τ* ^2^ = 1.

**Figure S13:**
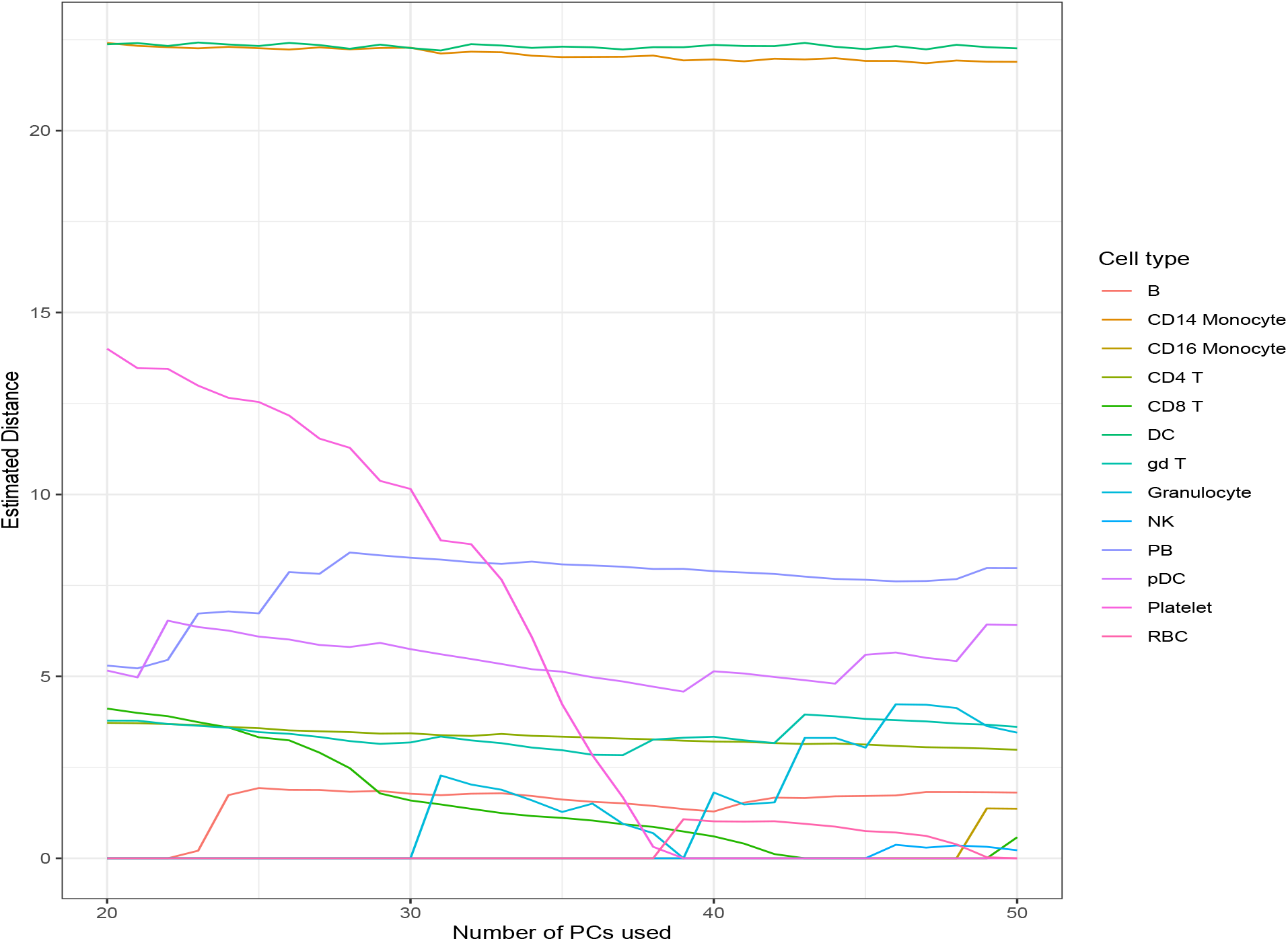
On the (1) dataset, distances were estimated using 20 ≥ *K* ≥ 50 PCs. For a majority of the cell types, the estimated distance is stable as *K* varies.

